# Cell-Permeable Morpholino Inhibits Programmed Death-Ligand 1 at mRNA Level and Potentiates Antitumor Immunity in Breast Cancer

**DOI:** 10.64898/2026.01.02.697345

**Authors:** Sudipta Sarkar, Ujjwal Ghosh, Subhamoy Pratihar, Surajit Sinha

**Affiliations:** Indian Association for the Cultivation of Science, School of Applied and Interdisciplinary Sciences, Kolkata 700032, India

**Keywords:** Cell permeable morpholino, Antisense therapy, Anti-PD-L1 therapy, tumor regression, Off-target effect, Durvalumab immunotherapy

## Abstract

Morpholino-based ASOs with nuclease-resistant backbone and high sequence specificity could modulate the PD-L1/PD-1 axis to overcome T regulatory cell mediated inhibition but poor cell-permeability limits their clinical applications. Here, we develop a guanidinium-modified cell-permeable morpholino specifically targeting PD-L1 in breast cancer. It suppresses PD-L1 expression through effectively mitigating cytokine induced PD-L1 upregulation and enhancing autophagic flux. Notably, the ASO synergistically with durvalumab (anti-PD-L1 antibody) disrupt, hyperglycosylated PD-L1 by mRNA-level interference and also reduced PD-1⁺ T-cell populations. Importantly, systemic and intra-tumoral administration of ASO demonstrated no abnormalities in cardiac tissues which show no off-target toxicity but it reprogrammed the tumor microenvironment by upregulating inducible nitric-oxide synthase and suppressed regulatory cells, collectively enhancing antitumor immune responses. Together, these findings establish morpholino ASO as a promising targeting therapeutic strategy to overcome tumor-induced immunosuppression, opening new avenues for precision treatment in breast-cancer.

## Introduction

Cancer immune evasion mechanism is the interplay involving Programmed Death-Ligand 1 (PD-L1) and its receptor, Programmed Death-1 (PD-1), both of which play pivotal roles in facilitating immune evasion by cancer cells leading to progression and metastasis of the disease. The inhibitory actions of PD-L1 on cancer cells and PD-1 on T cells serve as principal impediments, predominantly thwarting T cell-mediated cytotoxicity against tumour cells. Consequently, targeting this intricate interplay holds the promise of augmenting T cell-mediated cytotoxicity, rendering tumour cells more susceptible to conventional chemotherapy^1–4^. Over the past decade, antibody-based immune therapy has revolutionized the management of advanced-stage cancers, including lung cancer^5^, breast^6^, melanoma^7^, and other malignancies^8^. However, the efficacy of disrupting this interaction employing immune therapy yields a response rate ranging merely between 15-30%^1,9^. This suboptimal response is attributed to the intricate intracellular regulation of PD-L1, which remains inadequately elucidated. Moreover, the absence of antibodies capable of modulating the expansive array of this signalling cascade further complicates therapeutic interventions^9^. Hence, new approaches are under development for a broad range coverage of immune therapy^10,11^.

PD-L1, a member of the B7 family characterized as a type-I transmembrane glycoprotein^4^, undergoes upregulation in response to interferon-gamma (IFN-γ) stimulation and predominantly exerts its influence through the intricate PI3K-Akt and Stat3 pathways. Notably, GSK-3β emerges as a key regulator^12^, exerting post-transcriptional control over PD-L1, thereby aligning itself with the recognized hallmarks of cancer^13^. Recent research increasingly targets autophagy-driven tumour eradication as a promising anti-cancer strategy. Key regulators LC3 and Beclin are central to autophagy initiation, and elucidating their expression dynamics may optimize therapeutic approaches^14^. However, the precise influence of autophagy-mediated T cell responses to anti-cancer drugs remains unclear. Thus, we endeavour to ascertain whether targeting PD-L1 at the mRNA level by morpholino ASO can effectively modulate the aforementioned pathway to augment immunotherapeutic interventions.

In light of the recognized advantages and limitations of protein-centric therapeutics, the advent of antisense oligonucleotide (ASO) modalities targeting a broad repertoire of oncogenic transcripts has emerged as a vanguard in contemporary research, endeavoring to modulate these tumor-promoting genes while circumventing the evolution of resistance mechanisms^15^. In ASO research, phosphorodiamidate morpholino oligonucleotides (PMOs) have become a promising RNA-targeted agent as four morpholino based drugs (Eteplirsen, Golodirsen, Viltolarsen and Casimersen) have been approved by US FDA for the treatment of Duchenne Muscular Dystrophy (DMD)^16^ which open a new therapeutic regime targeting other genetic diseases. In the context of the challenges in developing immunotherapies, we present a novel approach involving a PD-L1 targeting, cell-permeable guanidinium-linked PMO (PD-L1 GMO-PMO chimera, termed PGP), which binds mRNA without requiring any delivery vehicles or transfection reagents. Unlike conventional immunotherapies, PGP ASO specifically targets PD-L1 at the mRNA level in breast cancer. Moreover, its small size (∼8 kDa) and nuclease stability suggest a simpler and more efficient therapeutic regimen compared to standard immunotherapy.

## Materials and Methods

### Solid phase synthesis of GMO-PMO and PMO

The GMO (Guanidinium Morpholino Oligonucleotides)-PMO (Phosphorodiamidate Morpholino Oligonucleotides) chimera and PMO were synthesized according to previously reported protocol^17, 18^ with ensuring its binding ability towards mature mRNA as shown in tables S1 & S2. After final trityl deprotection, the resin bound morpholino oligonucleotide was heated at 55°C in 30% aq-NH_3_ for 16 hrs. Then the aliquot was lyophilized and redissolved in water (100 µL) and subjected to acetone (1:7 v/v) precipitation. Purity was checked by HPLC and characterized by MALDI-TOF. The final compound was lyophilized to dry and dissolved to biological grade water for assay.

### Cells

Human triple-negative breast cancer cell lines MDA-MB-231, MDA-MB-468, and T-cell lymphoblast Jurkat E6.1 were obtained from NCCS, Pune and maintained in appropriate media: either 10% Fetal Bovine Serum (FBS)-MEM with non-essential amino acids, DMEM high glucose, or RPMI-1640, respectively, in T25 culture flasks, with Penicillin-Streptomycin-Glutamine (100X) (Gibco, USA). The 4T1 murine breast cancer cells (ATCC CRL-2539) were gifted by Dr. Saptak Banerjee, CNCI, Kolkata. The 4T1 cells were maintained in 10% bovine calf serum (Gibco)-RPMI under the same conditions.

### Immunoblot

Cells were seeded at 80% confluency in either 35mm culture dishes or 6-well plates, with the appropriate complete growth media. After 24 hours, various concentrations of PGP ASO were added to the media. After 48 hours of incubation, cells were lysed using RIPA buffer, PMSF (Sigma, 10837091001), and a phosphatase inhibitor cocktail (Sigma). Western blot analysis was performed using the following antibodies: anti-PD-L1 (13684, Cell Signalling, 1:1000), anti-JAK-2 (A19629, ABclonal, 1:1000), anti-IRF-1 (A7692, ABclonal, 1:1000), anti-FOXO-3A (A0102, ABclonal, 1:1000), anti-phospho-STAT-3-Y705 (AP0705, ABclonal, 1:1000), anti-phospho-Akt-1/2/3 (sc-514032, Santa Cruz, 1:1000), anti-P21 (A19094, ABclonal, 1:1000), anti-Cdc-42 (2462, Cell Signalling, 1:1000), anti-GSK-3β (27C10, Cell Signalling, 1:1000), anti-GAPDH (2118, Cell Signaling, 1:1000), anti-NF-κB (PA5-37718, Invitrogen, 1:1000), anti-Akt1 (A11016, ABclonal, 1:1000), anti-mTOR (A11354, ABclonal, 1:1000), anti-LC-3B I/II (PA1-16931, Invitrogen, 1:1000), anti-Beclin (MA5-32938, Invitrogen, 1:1000), anti-Bax (2772, Cell Signalling, 1:1000), anti-P62 (66184-1-IG, Proteintech, 1:1000), and anti-cleaved caspase 9 (9505, Cell Signalling, 1:1000).

### Ascites Carcinoma mediated allograft tumor Model

Balb/C albino mice (4-8 weeks of age) were obtained from the State Centre for Laboratory Animal Breeding, West Bengal Livestock Development Corporation Limited (a Government of West Bengal undertaking), Kalyani, West Bengal, with approval from the animal ethical committee (IACS/IAEC/S/2023/SS-02). Briefly, Ehrlich Ascites Carcinoma (EAC) cells were harvested intraperitoneally from the mice, followed by centrifugation at 200 g. The pellet was resuspended in RPMI-1640 medium and allografted subcutaneously with ≥90% cell viability. After 12 days, mice were administered mice specific PD-L1 GMOPMO antisense oligonucleotide (mPGP ASO) or scramble PDL-1 GMOPMO oligonucleotide (mPGP SCR) at a dose of 5 mg/kg body weight via intratumorally and intravenous injections, administered every 3 days. Intratumoral group is named as PD-L1 Knockdown-intratumoral (PD-L1 KD-IT) and the intravenous group was named as PD-L1 Knockdown-Intravenous (PD-L1 KD-IV). After 20 days, mice were euthanized, and tumours were excised. Blood samples were collected from the heart and portal vein, and other organs were harvested according to experimental requirements. Tumour volume was calculated using the formula: V (mm³) = (length × width²)/2.

### siRNA transfection

MDA-MB-231 or MDA-MB-468 cells were seeded onto 6-well plates at 80% confluency and subsequently transfected with PDL-1 siRNA (AM16708, Thermo Fisher) or negative control siRNA (AM4611, Thermo Fisher) using Lipofectamine™ 3000 Transfection Reagent (L3000001, Thermo Fisher). After 48 hours of incubation, the cells were either lysed with RIPA buffer or processed for immunocytochemistry analysis.

### Immunocytochemistry and Immunofluorescence

MDA-MB-231 cells were seeded onto poly-L-lysine-coated coverslips at 40% confluency. After 24 hours, treatments were applied, including morpholino oligos (PGP ASO, PGP SCR, mPGP ASO, mPGP SCR), siRNA targeting PD-L1. After a 48-hour incubation period, cells were fixed with 4% paraformaldehyde in PBS. Following fixation, cells were permeabilized using 0.1% Triton-X and blocked with 10% goat serum in 1% BSA-PBS. Cells were then incubated overnight at 4°C in a humidified chamber with primary antibodies against PD-L1 (13684, Cell Signaling, 1:200). After primary antibody incubation, cells were incubated with secondary antibodies (anti-rabbit Alexa Fluor 488, 1:200, or anti-mouse Alexa Fluor 594, 1:200) for 1 hour at room temperature. DAPI (1 μg/ml) was added for 5 minutes, and coverslips were mounted using Slowfade Gold (Invitrogen) and sealed. Images were captured using the Zeiss BIG.2 microscope at 63× magnification or the Leica TCS STED confocal system after 24 hours.

For ascites solid tumor analysis, slides were processed following deparaffinization (using graded ethanol), antigen unmasking (at pH 6), and permeabilization (with 0.25% Triton-X). After blocking, slides were incubated overnight at 4°C with primary antibodies against PD-L1 (A11273, ABclonal, 1:200; 29122, Cell Signalling, 1:200), LC3B (PA1-16931, Invitrogen, 1:200), Beclin (MA5-32938, Invitrogen, 1:200), Bcl2 (15071, 2876, Cell Signalling, 1:200), mTOR (A2445, ABclonal, 1:200), PD-1 (A11973, ABclonal, 1:200), and iNOS (MA5-17139, Invitrogen, 1:200). The next day, slides were incubated with secondary antibodies (anti-rabbit Alexa Fluor 488, 1:200, or anti-mouse Alexa Fluor 594, 1:200) for 1 hour at room temperature. Slides were then mounted with DAPI (1 μg/ml) and Vectashield for imaging. The images were captured using the Zeiss BIG.2 or Leica TCS STED confocal system.

### Morpholino transfection and lysosomal escape study

MDA-MB-231 cells were seeded onto dishes for 24 hours. PD-L1 GMO-PMO ASO (PGP ASO) and PD-L1 GMO-PMO SCR (PGP SCR) were conjugated with BODIPY at the 3′ end. The cells were then exposed to these oligonucleotides for 4 and 20 hours. Following incubation, Lysotracker Red DND-99 (L7528, Invitrogen) was added and incubated for an additional hour. Live cell imaging was subsequently performed using a Zeiss BIG.2 microscope.

To evaluate the transfection efficiency in MDA-MB-231 and MDA-MB-468 cells, BODIPY-conjugated oligonucleotides, including regular PMOs conjugated with BODIPY, were incubated for 4 hours. The cells were then processed using the BD FACS Aria III cytometer and analyzed with FlowJo software.

### IFN-γ induction analysis

Human Interferon-γ (hIFN-γ) Protein (10 ng/mL) (80385, Cell Signalling Technology) was added to 10% FBS-DMEM or MEM (NEAA) medium for 24 hours. Subsequently, PGP ASO and PGP SCR were introduced for an additional 48 hours. After the incubation period, cells were lysed with ice-cold RIPA buffer for further analysis.

### Endocytosis pathway analysis

MDA-MB-231 and MDA-MB-468 cells were seeded into 6-well plates at 70–80% confluency and incubated for 24 hours. PGP ASO (1 μM) and PGP SCR (1 μM) were then added for 48 hours in 10% FBS-DMEM or MEM (NEAA). Subsequently, pharmacological blockers—Amiloride (20 μM) (A7410, Sigma), Genistein (20 μM) (G6649, Sigma), Chlorpromazine (5 μM) (C8138, Sigma), Methyl-β-cyclodextrin (10 μM) (C4555, Sigma), and Chloroquine (10 μM) (C6628, Sigma)—were introduced for an additional 6 hours in 0.5% FBS-DMEM or MEM (NEAA) prior to cell harvest for Western blot analysis.

### Monoclonal antibody mediated assay

Durvalumab, a humanized recombinant monoclonal antibody (MA5-42048, Invitrogen), was used to block PD-L1 on the cell membrane. Briefly, MDA-MB-231 and MDA-MB-468 cells were seeded into 6-well plates and incubated for 24 hours. Durvalumab was then added at two concentrations (7.2 ng/mL and 14.4 ng/mL), either alone or in combination with PGP ASO (1 μM), and the cells were incubated for an additional 48 hours in full growth media (10% FBS-DMEM or MEM (NEAA)). After incubation, the cells were lysed using RIPA buffer, and immunoblotting was performed to analyses PD-L1 and related proteins.

### RT-qPCR

MDA-MB-231, MDA-MB-468, cells were seeded onto 6-well plates at 90% confluency and incubated for 24 hours. The following day, MDA-MB-231 and MDA-MB-468 cells were treated with PGP ASO (1 μM), PGP SCR (1 μM), and durvalumab (7.2 ng/mL) in 10% FBS-DMEM or MEM (NEAA) media for 48 hours. After the incubation period, cells were lysed using TRIzol™ Reagent (15596026, Invitrogen) and processed with the total RNA Miniprep Kit (T2010S, New England Biolabs), following the manufacturer’s protocol.

cDNA was synthesized from 1–5 μg of RNA using the RevertAid First Strand cDNA Synthesis Kit (K1621, Thermo Scientific). For analysis, 2 μl of cDNA was used for agarose gel electrophoresis, and 100–500 ng of cDNA was used for qPCR analysis with PowerUp™ SYBR™ Green Master Mix for qPCR (A25741, Applied Biosystems), following the manufacturer’s protocol. All primers are listed in table S3.

### Cell membrane PD-L1 analysis

MDA-MB-231 cells were seeded onto confocal dishes at 50% confluency. Following treatment with PGP ASO and PGP SCR, either alone or in combination with durvalumab (7.2 ng/mL and 14.4 ng/mL) and IFN-γ (10 ng/mL), cells were washed with ice-cold 1× PBS and subsequently fixed with 4% PFA-PBS.

For immunostaining, cells were incubated overnight with a primary antibody targeting the extracellular domain of PD-L1 (D8T4X, 86744, Cell Signalling Technology; 1:200), followed by blocking with 10% goat serum in 1% BSA-PBS. The next day, cells were incubated with an Alexa Fluor 488-conjugated anti-rabbit secondary antibody (1:200) for 1 hour. Finally, samples were mounted using SlowFade Gold (Invitrogen) containing DAPI. Imaging was performed using a Zeiss BIG.2 microscope at 63× magnification, and image processing was conducted with LAS X software.

### Cell fractionation

MDA-MB-231 cells were seeded onto 100 mm culture dishes at 90% confluency and incubated for 24 hours. The following day, PGP ASO, PGP SCR, and durvalumab (7.2 ng/mL) were added as previously described. After incubation, the cells were washed with ice-cold 1× PBS and subsequently lysed using a cell fractionation buffer.

For cytosolic and nuclear fractionation, cells were gently collected by scraping, pelleted by centrifugation, and resuspended in a hypotonic buffer solution (20 mM Tris-HCl, pH 7.4, 10 mM NaCl, 3 mM MgCl₂). The suspension was incubated for 15–30 minutes with intermittent vortexing. Following centrifugation, the supernatant, representing the cytosolic fraction, was separated, while the pellet was further processed for nuclear extraction.

The cell pellet was resuspended in nuclear extraction buffer (10 mM Tris-HCl, pH 7.4, 2 mM Na₃VO₄, 100 mM NaCl, 1% Triton X-100, 1 mM EDTA, 10% glycerol, 1 mM EGTA, 0.1% SDS, 1 mM NaF, 0.5% sodium deoxycholate, and 20 mM Na₄P₂O₇) and incubated for 30 minutes to 1 hour with intermittent vortexing. Following centrifugation, the supernatant containing the nuclear extract was collected. A total of 30–50 μg of protein was used as input for immunoblot analysis.

### FESEM

Cell suspensions were cultured on poly-L-lysine-coated coverslips for 24 to 48 hours, contingent upon the specific adherence properties of each cell type. Following successful attachment, cells were subjected to various drug treatments, including PGP ASO. After a 48h incubation period, cells were fixed with 2.5% glutaraldehyde, followed by secondary fixation with osmium tetroxide for 1 hour each. Subsequently, the specimens underwent a graded ethanol dehydration series, followed by HMDS drying. The prepared samples were then mounted, sputter-coated, and imaged using a JEOL Field Emission Scanning Electron Microscope (FESEM).

### Co-culture with Jurkat E6.1

Jurkat E6.1 cells were cultured in RPMI-1640 medium supplemented with 10% FBS. For co-culture experiments, Jurkat E6.1 cells and triple-negative breast cancer (TNBC) cell line (MDA-MB-231) were co-cultured at a 6:1 ratio. Co-cultures were established through a trans-well co-culture system (Cell Culture Insert, Cell Bio, TCP241) for Western blot and flow cytometry analyses.

Jurkat E6.1 cells were pre-activated with Cell Activation Cocktail (423301, BioLegend) for 14 hours, followed by centrifugation and washing with 1× PBS. The activated Jurkat cells were then co-cultured with pretreated TNBC cells for 24 hours. After incubation, the lower fraction (TNBC cells) was washed and lysed using RIPA buffer for immunoblot analysis. The upper fraction (Jurkat E6.1 cells) was pelleted, washed, fixed with paraformaldehyde (PFA), and incubated with a primary PD-1 antibody, followed by blocking and staining with the respective secondary antibody. Finally, flow cytometric analysis was performed using the BD FACS Aria III cytometer and FlowJo software.

### ELISA

Blood was collected from allograft tumor bearing mice into sterile microcentrifuge tubes. Serum was isolated by centrifuging the whole blood at 2000 × g and stored at −20°C until further analysis. Enzyme-linked immunosorbent assays (ELISA) for IFN-γ (ELM-IFNg-1, RayBiotech) and TNF-α (ELM-TNFa-1, RayBiotech) were performed according to the manufacturer’s protocol.

### Flow cytometric analysis of Single cell suspension

Ascites tumors were isolated and meticulously dissected into 1 mm³ fragments using a sterile scalpel. The tissue fragments were enzymatically digested in a digestion buffer composed of collagenase (150,000 U), DNase I (1 mg/mL), and 5% FBS-RPMI-1640 at 37°C for 30 minutes. The resulting cell suspension was filtered through a 40 μm sterile cell strainer (TCP024, Cell Bio), washed with 2% FBS-PBS containing EDTA, and centrifuged at 300 × g to obtain a cell pellet. Erythrocyte lysis was performed using ammonium chloride buffer, and the final cell pellet was resuspended in an appropriate medium for subsequent analyses.

For live-cell analysis, cells were maintained in 10% FBS-RPMI-1640 medium at 37°C in a 5% CO₂ humidified incubator until further processing. For cell marker analysis, cells were fixed with 4% PFA-PBS, permeabilized using a 0.1% Triton-X solution for intracellular staining, and subsequently incubated with fluorophore-conjugated primary antibodies against FOXP3 (563101, BD Biosciences; 1:200), CD4 (12-0041-82, eBioscience™; 1:200), CD8 (MHCD0804, Invitrogen; 553031, BD Biosciences; 1:200), and PD-1 (A11973, ABclonal; 1:200). Blocking was performed using 1% BSA-PBS to minimize non-specific binding.

### Intra-tumoral ROS generation

Single-cell suspensions were prepared as previously described. Cells were then incubated with DCFDA (10 μM) for 30 minutes at 37°C, followed by washing with 1× PBS and resuspension in flow cytometry buffer. Subsequent analysis was conducted using the BD FACS Aria III cytometer and FlowJo software.

### Intra-tumoral mitochondrial potential analysis

Tumors from different treatment groups, including PD-L1 Knock down mice and the corresponding SCR group, were isolated and processed to generate single-cell suspensions. These cells were then incubated with JC-1 (T3168, Invitrogen) using valinomycin as a positive control, following the manufacturer’s protocol.

### Intra-tumoral BrdU incorporation assay

Single cells isolated from ascites tumors were processed and subsequently incubated with BrdUTP. Apoptotic analysis was performed using the APO-BrdU™ TUNEL Assay Kit, with Anti-BrdU Alexa Fluor™ 488 (A23210, Invitrogen), following the manufacturer’s protocol.

### Caspase Glo-9 assay

MDA-MB-231 and MDA-MB-468 cells were seeded into 96-well plates at 30% confluency and incubated for 24 hours. The following day, PGP ASO, PGP SCR, and Durvalumab, either alone or in combination with PGP ASO, were added and incubated for 48 hours. Following incubation, the cells were lysed, and luminescence was measured according to the manufacturer’s protocol using the SpectraMax ID5 Multi-Mode Microplate Reader Platform.

### Serum toxicity assessment

Blood was collected via intracardiac syringe extraction and allowed to clot before being centrifuged at 3,000 rpm for 20 minutes to obtain serum. The serum was then used for biochemical assays, including SGOT, SGPT, albumin, serum urea, creatinine, and CK-MB activity. These assays were performed using Transasia ERBA commercial kits, following the manufacturer’s protocol.

### H&E staining

Ascites tumor-bearing mice were sacrificed, and their organs were harvested, washed with PBS, and stored in 10% neutral-buffered formalin. The tissues were then prepared for microtome sectioning, followed by staining with hematoxylin and counterstaining with eosin. After staining, the tissue sections were mounted with DPX, and images were captured using an inverted fluorescence microscope (Olympus IX51) and a Leica fluorescence microscope.

### Statistical Analysis

All data are reported as the mean ± SEM. GraphPad Prism software Version 8 was used for the generation of all bar graphs and statistical analyses. Statistical analyses were performed with one or two-way ANOVA and student t test for multiple comparisons between different treatment groups across different assays as specified in the respective legends.

## Results

### PD-L1 GMO-PMO suppressed targeted proteins in a dose dependent manner in both TNBC cell lines

We chose triple-negative breast cancer (TNBC), marked by its aggressive phenotype and lack of ER, PR, and HER2, remains the most chemo-resistant and therapeutically unresponsive breast cancer subtype. Its poor prognosis stems from limited sensitivity to endocrine or HER2-targeted regimens^19–21^. In prior work, we demonstrated that unlike regular PMO, GMO-PMO chimera (for structure **Figure S1**) exhibits enhanced cellular uptake, nuclease resistance, aqueous solubility and low cytotoxicity. This chimera effectively inhibited NANOG and potentiated Taxol response^22^. As part of our ongoing research toward the applications of GMO-PMO chimera, here we reported the anti-cancer potential of GMO-PMO chimera (PD-L1 GMOPMO; PGP) (**Table 1, For Blast ID and FASTA format of PD-L1 Tables S1 and S2)** targeting PD-L1 (an immune-suppressive ligand of PD-1) at the mRNA level.

**Table 1:**
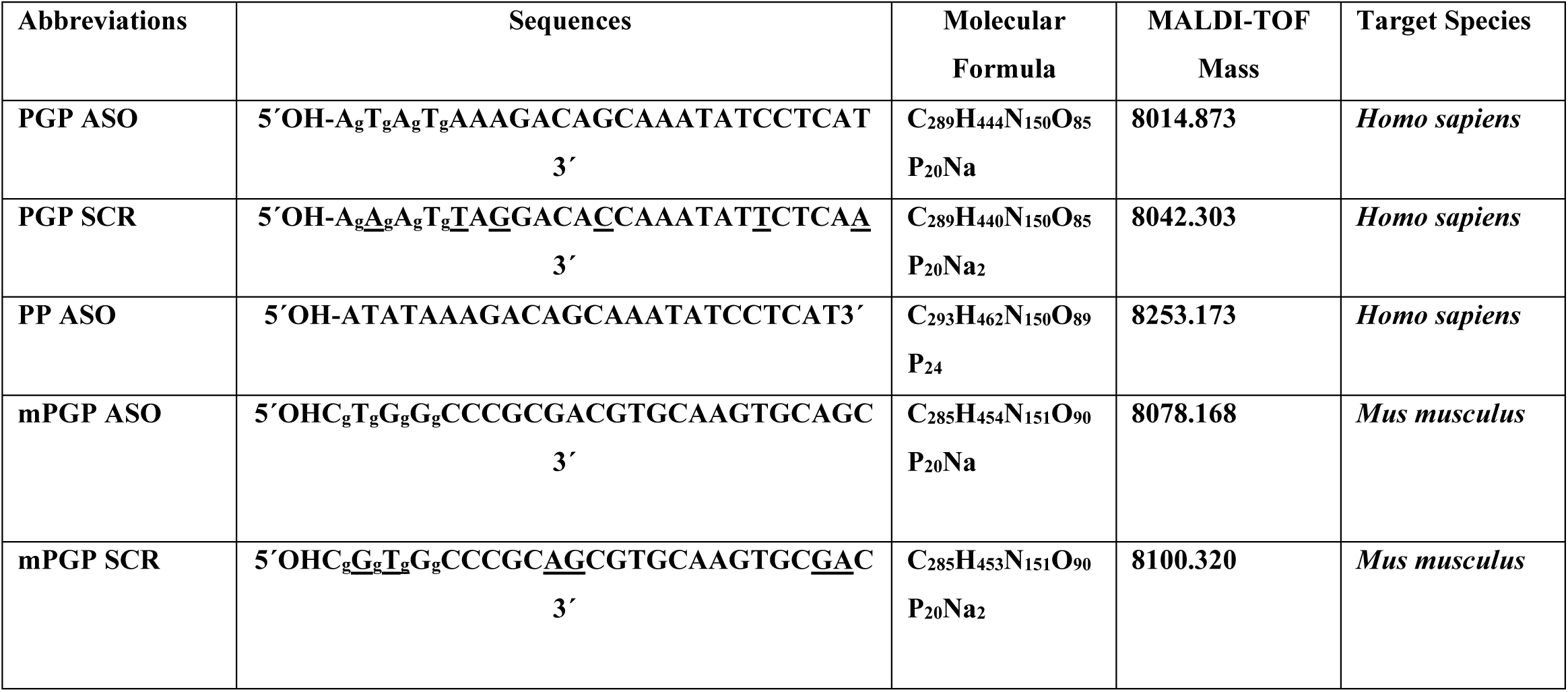
Morpholino Sequences and targeted species (g denotes guanidium linkages into the morpholino backbone, underline denotes the mismatch base in scramble sequence)

Before progressing into mechanistic study of PGP ASO *in vitro* and *in vivo*, first we purified and characterized the oligos by HPLC and MALDI TOF mass analysis, respectively (**Figure S2-6**) followed by the measurement of duplex stability by thermal melting (Tm) study with the complementary RNA strand. We observed that Tm of PP ASO: RNA, PGP ASO: RNA and PGP SCR: RNA was 55°C, 50°C and 41°C, respectively (**Table S4, Figure S7**). Though the Tm of PP ASO (stands for regular PMO) was 5°C higher than PGP ASO but it was ineffective in gene silencing due to the poor cell-permeable property (*vide infra*). In the case of PGP SCR, the duplex was destabilized by 14°C and was the reason for its inactivity.

To assess the antisense efficacy of PGP in TNBC cell lines (MDA-MB-231 and MDA-MB-468), a dose-dependent analysis was performed. PGP ASO elicited robust suppression of PD-L1 expression in both cell lines (**Figure 1A, B1, 2 & S8A**), accompanied by marked downregulation of its effectors JAK2 (0.329±0.045 at 1μM; 0.386±0.052 at 1.5 μM) and IRF-1(0.319±0.044 at 1μM; 0.181±0.012 at 1.5 μM). In contrast, scramble control (PGP-SCR) and non-permeable PP-PMO (regular PDL-1 PMO has no cell transfection properties) showed no inhibitory activity, affirming the sequence-specific action of PGP. Notably, FOXO3A, a negative regulator of PD-L1, was significantly upregulated in a dose-responsive manner (1.420±0.101 at 0.5μM; 1.238±0.037 at 1 μM; 1.526±0.066 at 1.5 μM) (**Figure 1 B1, 2**). Key oncogenic mediators of the Akt-Stat3 axis were concurrently suppressed, including phosphorylated Akt and Stat3. Downstream targets implicated in cell cycle regulation, such as Cdc42 and p21^Cip1^, were also declined in MDA-MB-231 cells, indicating disruption of proliferative signaling. Given p21^Cip1^’s transcriptional regulation by Stat3 and role in G1/S transition, its repression underscores PGP ASO’s antiproliferative capacity (**Figure S8B**). These molecular effects were further substantiated via PD-L1 immunocytochemistry (**Figure S8C)**. Collectively, the data highlight PGP ASO as a potent, mechanistically precise anti-cancer agent. Here, we mainly focused on MDA-MB-231 TNBC cell line which has low claudin expression, makes it more aggressive and resistance in chemotherapy to develop an anti-cancer drug. PGP ASO showed an inhibition from 1μM dose, hence was chosen for further studies.

**Figure 1:**
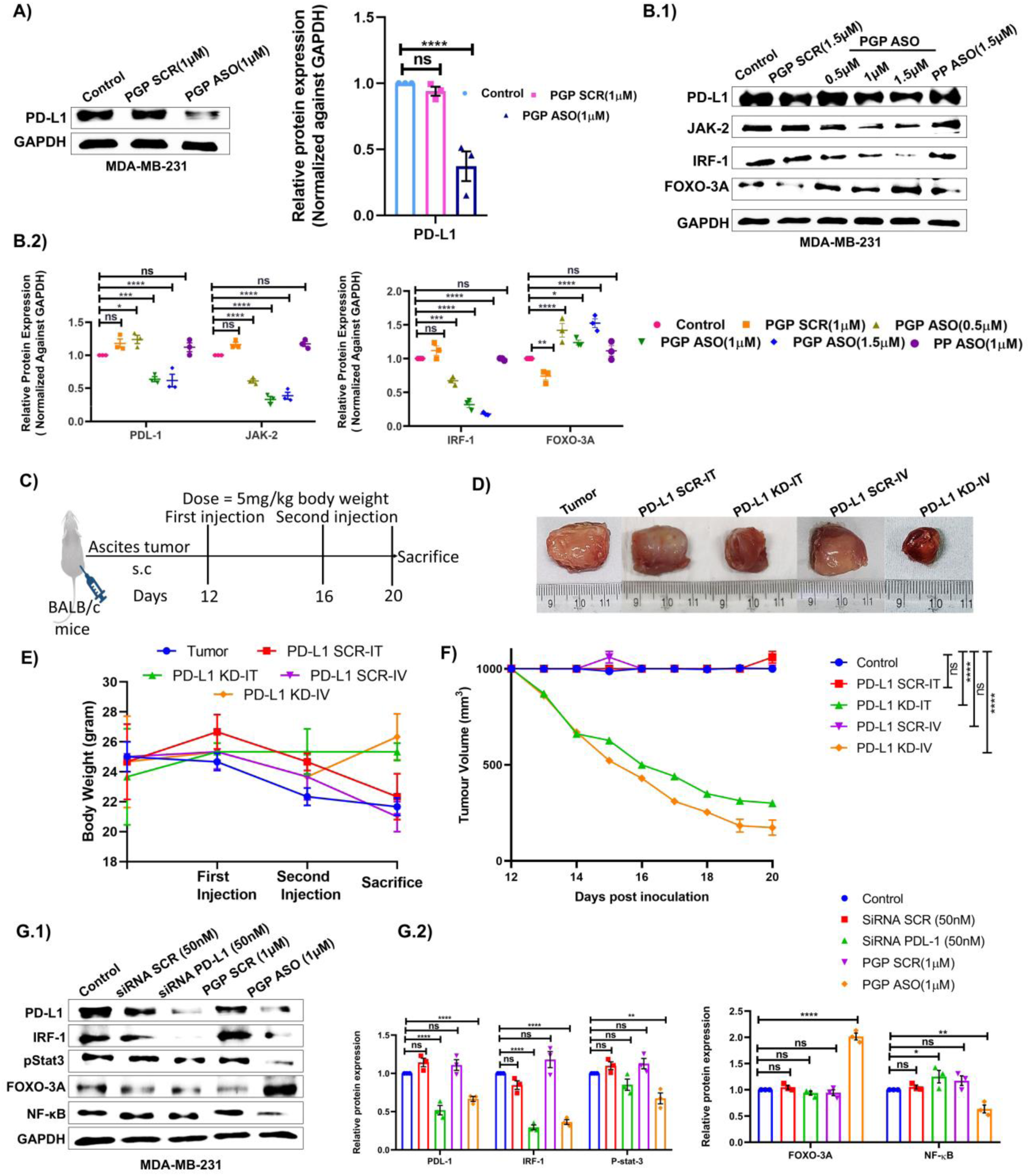
PD-L1 targeted inhibition by cell permeable GMO-PMO. **(A)**PD-L1 inhibition in MDA-MB-231 cell line (left panel) and bar diagram representation of PD-L1 inhibition (right panel). **(B.1 & B.2)** Representative images of western blot analysis of PD-L1 and associated protein in a dose dependent study of PGP ASO in MDA-MB-231 cells. And Graphical representation of dose dependent inhibition of PD-L1 and associated proteins in MDA-MB-231 cells. **(C)** Timeline of ascites allograft model generation, mPGP ASO and mPGP SCR treatments. **(D)** Representative images of ascites tumor after treatment duration. **(E)** Graphical representation of body weight measurement throughout the treatment period. **(F)** Graphical representation of tumor volume measurement throughout the treatment period. **(G.1 & G.2)** Bar diagram representation of densitometric analysis of PGP ASO and siRNA mediated comparative inhibition profiles and representative images of western blot analysis of PGP ASO and siRNA mediated comparative inhibition of PD-L1 and associated proteins. Data shown as mean ± SEM, Ordinary Two-way ANOVA was performed followed by Tukey multiple comparison test. \**p* ≤ 0.5 considered as statistically significant, ** and *** considered as higher significant than *p* ≤ 0.5.

Next, we developed Ehrlich Ascites Carcinoma (EAC) induced allograft tumor model, a spontaneous murine mammary adenocarcinoma^23^ (EAC cells specifically grown inside peritoneal cavity of mice, which makes it more aggressive tumor model) to extrapolate the PGP ASO mediated inhibition *in vivo*. Here we used *Mus musculus* specific PD-L1 GMO-PMO ASO (mPGP ASO, **Table 1**) via intratumorally and intravenous (5 mg/Kg body weight) for 7 days to knock-down PD-L1 (**Figure 1C)**. Based on whole tumor mass western blot analysis, a better effect in PD-L1 downregulation was observed when oligos were injected intratumorally (0.586 ± 0.055) than intravenously (0.707 ± 0.050) (**Figure 1 D-F)**. This observation was also confirmed through immunofluorescence of PD-L1 of tumor tissue (**Figure S8D**). JAK-2, pStat-3, IRF-1 all the proteins downregulated in both modes of injections, but pAkt 1/2/3 inhibited only at intravenous injection (**Figure S8E**).

### PGP ASO surpassed siRNA mediated inhibition of wide array of protein functions

The anti-cancer potential of exosome loaded siRNA-PDL-1^24^, PDL-1 theranostic^25^ and aptamer-ASO^26^ was reported in downregulation of PD-L1 and evaluated their therapeutic efficacy in cancer models. Though siRNA and morpholino worked in a different mechanism, morpholino have certain advantages over siRNA such as neutral backbone and nuclease stability. Furthermore, our modified morpholino (PGP) has a cell permeable property where no transfection reagents are required for its delivery. Unlike siRNA, PMOs were not easily accessible because their synthesis was challenging. Recently, the last hurdle for their synthesis by an automated oligo synthesizer has also been achieved by our group^27^. To evaluate the potential of our technology, we became interested to compare our PGP ASO with the commercially available siRNA against PD-L1. As expected, both PGP ASO and siRNA downregulated PD-L1 and their inhibition was 0.520±0.062 and 0.666±0.033 for siRNA and PGP ASO, respectively. Interestingly, PGP ASO showed better inhibitory activity on the downstream proteins of PD-L1 than siRNA with the expression of pStat-3 (0.851±0.074 for siRNA and 0.673±0.072 for PGP ASO) and NF-κB (1.252±0.118 for siRNA and 0.636±0.074 for PGP ASO) (**Figure 1 G1-2**). Furthermore, in MDA-MB-468 cell line IRF-1 was not inhibited by siRNA (**Figure S9**) and FOXO3A was only up-regulated in PGP ASO treated group (0.938±0.038 for siRNA and 2.016±0.064 for PGP ASO) (**Figure 1 G1-2 & S9**). Thus, PGP ASO not only effectively suppressed the expression of its intended target protein but also demonstrated the potential to modulate the activity of additional cancer-associated marker proteins. These findings underscore the therapeutic relevance of PD-L1-targeted inhibition in diverse subtypes of triple negative breast cancer.

### ASO-mediated inhibition of PD-L1 is marked by the lysosomal escape of PGP ASO

One of the most common challenges in the design and synthesis of antisense oligonucleotides (ASOs) is achieving efficient endosomal escape of the trapped oligonucleotides. To confirm that the inhibition of PD-L1 and associated proteins was not due to stress-induced effects, we conjugated Bodipy with PGP ASO and PGP SCR to evaluate the transfection efficiency of these oligonucleotides in MDA-MB-231 and MDA-MB-468 cell lines (**Figure S10 & 11A**). The results demonstrated approximately 50% transfection at 0.5 μM concentration (48.80 ± 1.20% for PGP SCR and 58.55 ± 1.45% for PGP ASO) and around 90% transfection at a 1 μM concentration (88.90 ± 2.10% for PGP SCR and 90.45 ± 1.35% for PGP ASO), while PP ASO and Bodipy fluorophore itself exhibited no transfection. These findings further confirmed that the guanidinium group enhances the cellular permeability of oligonucleotides, and that both scramble and antisense sequences exhibited similar transfection efficiencies (**Figure S11B**).

To investigate the potential lysosomal entrapment of oligonucleotides, cells were incubated with bodipy-labelled oligonucleotides and subsequently stained with lysotracker dye. The Pearson’s overlap coefficient decreased from 0.8 at 4 hours to 0.4 at 20 hours, indicating that both PGP ASO and PGP SCR successfully escaped from the lysosomes within 20 hours (**Figure S11C & D**).

### PGP ASO effectively abrogated IFN-γ induced PD-L1 expression

To investigate cytokine-induced PD-L1 expression, we used IFN-γ (Type II interferon) to mimic JAK-2/Stat-3/IRF-1 pathway upregulation in MDA-MB-231 and MDA-MB-468 cell line. In both cell lines pStat-3 decreased after PGP ASO treatment (0.591±0.017, 40.9% for MDA-MB-231 and 0.584±0.040, 41.6% for MDA-MB-468). IFN-γ secreted from tumor microenvironment showed dual effect on tumor migration and proliferation^28^. For the dual role of IFN-γ, opposite expression profile of Akt and mTOR in IFN-γ stimulated cell was observed (**Figure 2A, B & S12A, B**). Upregulation of LC-3B (one of major autophagy markers) was observed after co-treatment of IFN-γ and PGP ASO (2.708±0.104, 170.8% for MDA-MB-231 and 0.637±0.038 for MDA-MB-468), which led to autophagic progression of cancer cell. There was a prominent increase of NF-κB expression after IFN-γ treatment. Interestingly, Beclin was upregulated in MDA-MB-468 (66% for IFN-γ and PGP ASO co-treatment group and 29.6% for PGP ASO alone group) but downregulated in MDA-MB-231 upon PGP ASO treatment with (35.9%) or without IFN-γ stimulation (45%) (**Figure 2C, D & S12C, D**). In all the cases whether IFN-γ showed a positive or negative effect on protein expression, PGP ASO always inhibited the target protein. In *in vivo* scenario, both Beclin and LC3B level were increased in tumor section as shown in **Figure 2E & S13A**. When we studied this phenomenon in PD-L1 KD mice, mTOR (a major regulator of autophagy) decreased along with Bcl2 (**Figure 2F & S13B**). The above-mentioned data was also corroborated well with immunofluorescence study (**Figure 2G1-2),** where intravenous administration of mPGP ASO showed better efficacy. Therefore, even within the dynamic tumor microenvironment, where signaling pathways may undergo alterations, our synthesized PGP ASO is expected to retain its therapeutic efficacy.

**Figure 2:**
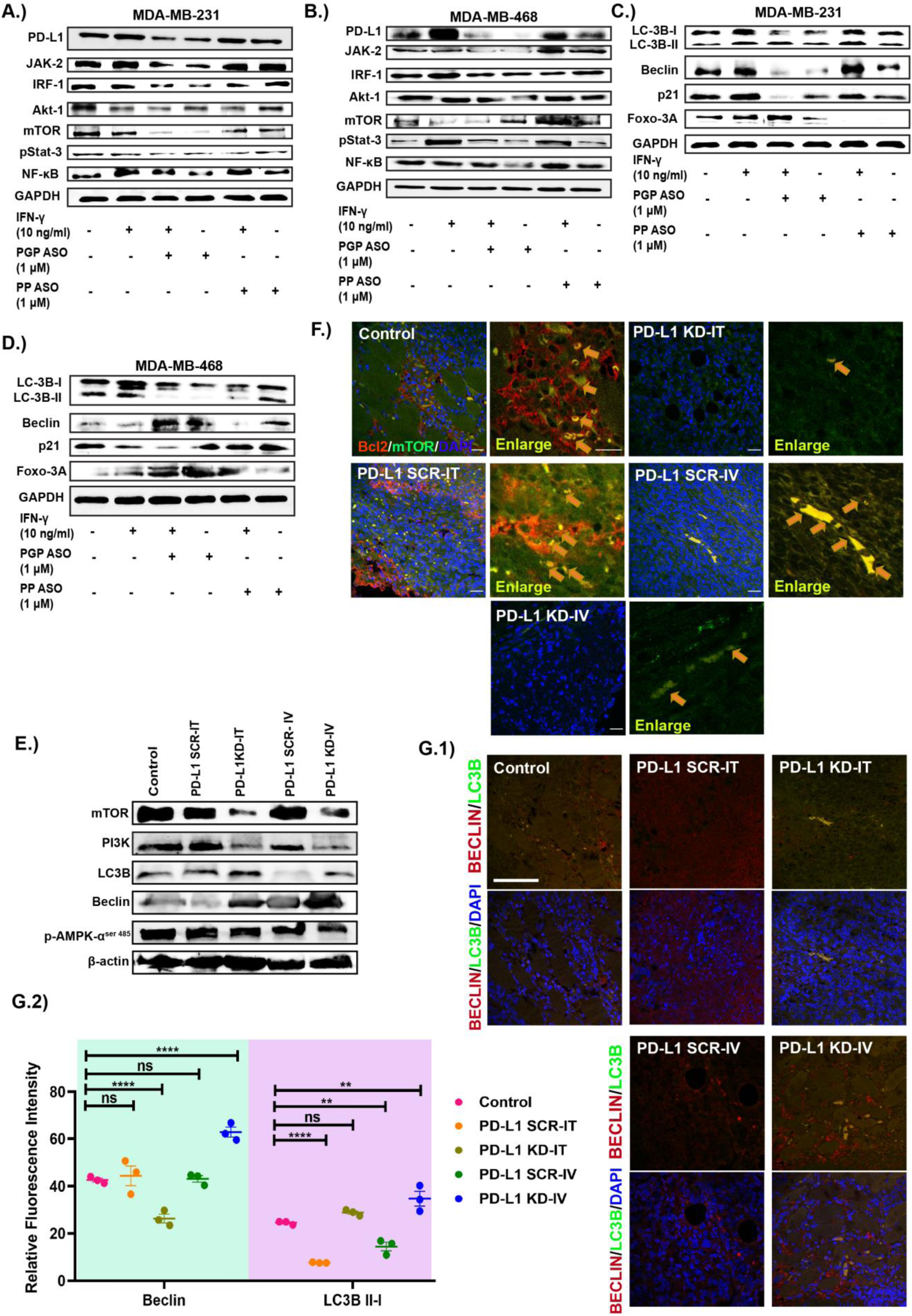
IFN-γ induced PD-L1 expression inhibited by PGP ASO. **(A)** PD-L1 and associated proteins expression pattern in MDA-MB-231 cell. **(B)** PD-L1 and associated proteins expression pattern in MDA-MB-468 cell**. (C)**Representative expression patterns of IFN-γ induced LC3B, Beclin, P21, FOXO-3A proteins in MDA-MB-231. **(D)**Representative expression patterns of IFN-γ induced LC3B, Beclin, P21, FOXO-3A proteins in MDA-MB-468**. (E)**Western blot analysis of mTOR and associated protein in PD-L1 KD/SCR IT/IV groups tumor tissue. **(F)** Immunofluorescence images of tumor tissue of different groups stained with Bcl2(red), mTOR(green) and DAPI (blue). Scale bar=5 μm, In enlarge figure Scale bar=10 μm. **(G.1 & G.2)** Immunofluorescence images of tumor section of different treatment groups stained with Beclin(red), LC3B (green) and counterstained with DAPI (Blue). Scale Bar = 20μm and relative fluorescence Intensity of Beclin and LC3B of G1 image. Data shown as mean ± SEM, Ordinary Two-way ANOVA was performed followed by sidak multiple comparison test. \**p* ≤ 0.5 considered as statistically significant, ** and *** considered as higher significant than *p* ≤ 0.5.

To investigate whether aforementioned PGP ASO mediated autophagy induction in MDA-MB-231 cells involves autophagosomal fusion with lysosome or not, we used chloroquine (hydroxy chloroquine; HCQ) to block lysosomal fusion with autophagosome^29,30^. First, we treated cells with chloroquine to check PD-L1 inhibition followed by PGP ASO incubation. Interestingly, HCQ and PGP ASO dual treated groups showed no PD-L1 inhibition though they have shown inhibition in single treated group which indicates PGP ASO might fused cellular proteins with lysosome to initiate autophagy. When lysosome fusion was blocked by the addition of HCQ, then PGP ASO failed to work. HCQ contradicts the PGP ASO mechanism. Similar trend was followed by PI3K, but the FOXO3A expression pattern indicates PGP ASO mediated upregulation was independent of lysosomal fusions. LC3B was upregulated in both HCQ as well as PGP ASO treatment. But LC3B is more punctate in HCQ than PGP ASO treatment, indicating accumulation of LC3B after HCQ treatment. HCQ and PGP ASO cotreatment opposed each other and LC3B level decreased to basal level (**Figure S14**).

### Intracellular shuttling of PD-L1 depends on endocytosis

PD-L1 nucleocytoplasmic trafficking is tightly governed by cytoskeletal elements. As shown in **Figure S8C**, nuclear PD-L1 localization persisted across groups, including those treated with PGP ASO, highlighting the nuclear delivery inefficiency of guanidinium-conjugated morpholino oligomers.

Glycosylated PD-L1 undergoes dynamic membrane-to-nuclear trafficking, maintaining cytosolic pools and repressing GSK-3β phosphorylation^31^. To assess trafficking dependent PD-L1 regulation, cells were exposed to endocytic inhibitors. Amiloride, genistein, and Mβ-CD modestly reduced PD-L1 relative to PGP ASO, whereas CPZ and chloroquine mirrored PGP ASO’s inhibitory profile in MDA-MB-231. In MDA-MB-468, only Mβ-CD suppressed PD-L1 (**Figure 3A**), indicating subtype-specific trafficking. PGP ASO induces autophagy, while chloroquine antagonistically reduces PD-L1 via autophagy inhibition, as discussed earlier in **Figure S14**.

**Figure 3:**
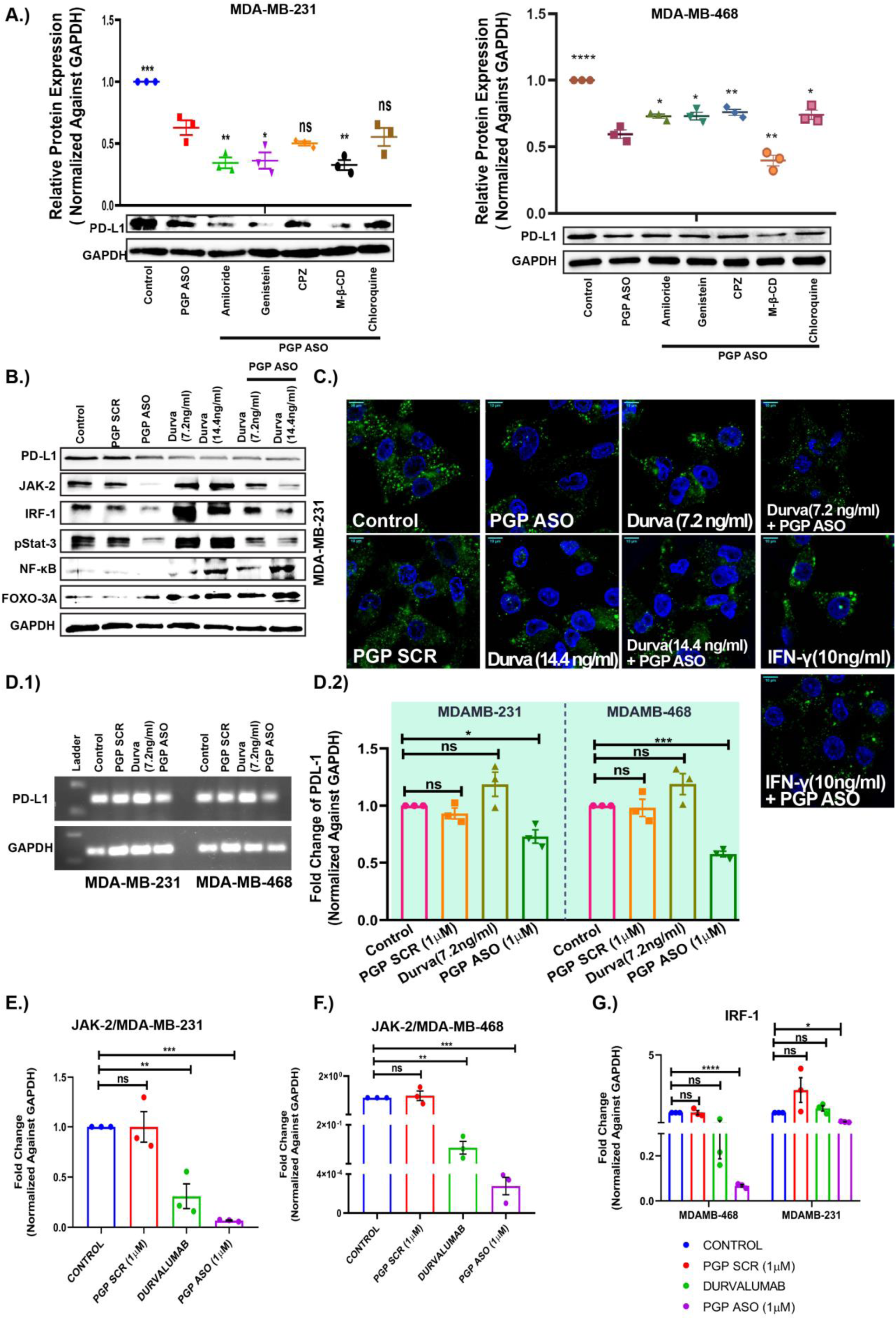
Cell membranous PD-L1 downregulated by PGP ASO mediated PD-L1 mRNA target. **(A)** Intracellular trafficking of PD-L1 in MDA-MB-231 and MDA-MB-468 cell line respectively. **(B)** Total protein analysis of PD-L1 and associated proteins after PGP ASO and durvalumab treatment in MDA-MB-231 cell. **(C)** Confocal images of Cell surface PD-L1 after different treatment conditions along with PGP ASO. Scale bar= 10μm. **(D.1 & D.2)** mRNA expression of PD-L1 after different treatment (Left panel) and graphical representation of densitometric analysis of PD-L1 mRNA expression (Right panel). **(E & F)** JAK-2 mRNA expression profile in MDA-MB-231 and MDA-MB-468 cell after different treatments respectively. **(G)** IRF-1 mRNA expression profile in MDA-MB-468 and MDA-MB-231 cells. Data shown as mean ± SEM, Ordinary Two-way ANOVA was performed followed by sidak multiple comparison test. \**p* ≤ 0.5 considered as statistically significant, ** and *** considered as higher significant than *p* ≤ 0.5.

### PGP ASO mediated mRNA blockade chemo-sensitize Glycosylated Surface PD-L1 inhibition by durvalumab

N-linked glycosylation stabilizes PD-L1 by preventing ubiquitin-proteasomal degradation. To assess whether glycosylated PD-L1 can evade mRNA-level suppression and restore cytosolic pools, durvalumab (a membrane PD-L1 targeting antibody drug) was applied alone and with PGP ASO. This strategy aimed to evaluate effects on total and membrane bound PD-L1, along with key signaling intermediates.

At 7.2 ng/mL, durvalumab reduced total PD-L1 to levels comparable with 1 μM PGP ASO. However, dual treatment did not further suppress PD-L1, implying a saturation point or compensatory stabilization of residual protein (**Figure 3B & S15A**).

Monotherapy with durvalumab markedly elevated JAK2, IRF-1, and pStat3 levels, whereas co-administration with PGP ASO mitigated their expression, indicating pathway convergence under PD-L1 inhibition. NF-κB remained persistently upregulated across all antibody-treated cohorts, reflecting sustained pro-inflammatory signaling. Notably, FOXO3A was robustly induced by PGP ASO, with maximal enhancement under 14.4 ng/mL durvalumab co-treatment, implicating PD-L1 in its post-translational suppression. Dual blockade at the transcript and membrane levels synergistically reinstated FOXO3A activity (**Figure 3B**). Immunogenic mediator induction likely stems from ADCC, as durvalumab (IgG1) facilitates Fc-dependent cytotoxicity^32^. Besides this, MDA-MB-468 also showed similar expression patterns (**Figure S15 B1,2**).

Surface PD-L1 intensity was comparable across 7.2 ng/mL, 14.4 ng/mL durvalumab, and PGP ASO monotherapy; notably, the 7.2 ng/mL combination mirrored PGP ASO alone in surface PD-L1 expression. (**Figure 3C & S15C**). Intriguingly, IFN-γ did not elicit cell membrane PD-L1 expression.

Furthermore, mRNA level of PD-L1 was reduced only after PGP ASO mediated mRNA target but remain unaltered in case of durvalumab, clearly showing surface PD-L1 blocking strategy is less effective than mRNA level manipulation (**Figure 3 D1,2**). Similarly, JAK-2 and IRF-1 mRNA levels were also significantly inhibited in PGP ASO group in both TNBC cell lines (**Figure 3E-G**).

### Autophagosome activation was pertinent to PGP ASO treatment, both in isolation and in conjunction with Durvalumab co-administration

PGP ASO monotherapy upregulated LC3B and Beclin, with Beclin elevated by 280% under co-treatment with 14.4 ng/mL Durvalumab, while LC3B induction remained exclusive to PGP ASO alone (**Figure S15D**). These findings suggest that translational silencing of PD-L1 via PGP ASO facilitates autophagy initiation. Although PGP ASO alone activates autophagy, dual inhibition—translational and membranous—may potentiate therapeutic efficacy.

Thus, PGP ASO-mediated PDL-1 silencing confers dual inhibition of oncogenic signaling and immune escape.

### PGP ASO inhibits nuclear transportation of PD-L1 and associated proteins

Nuclear PD-L1 may create a resistance towards immunotherapy. To investigate whether PGP ASO can overcome this phenomenon or not, we fractionated MDA-MB-231 cells to cytosol and nuclear fraction. In the case of PD-L1, PGP ASO and durvalumab showed downregulation in both fractions. But PGP SCR showed a prominent expression of PD-L1 in the nucleus fraction which indicated a feedback mechanism of cancer cells upon morpholino treatment. In the case of PGP ASO, it blocks mRNA in cytosol, downregulates PD-L1 nuclear transport as well as cytosolic presence. But PGP SCR lacks the inhibitory effect which concomitantly increased nuclear PD-L1 expression (**Figure 4 A1,2**). This observation further proves targeted inhibition of PGP ASO.

**Figure 4:**
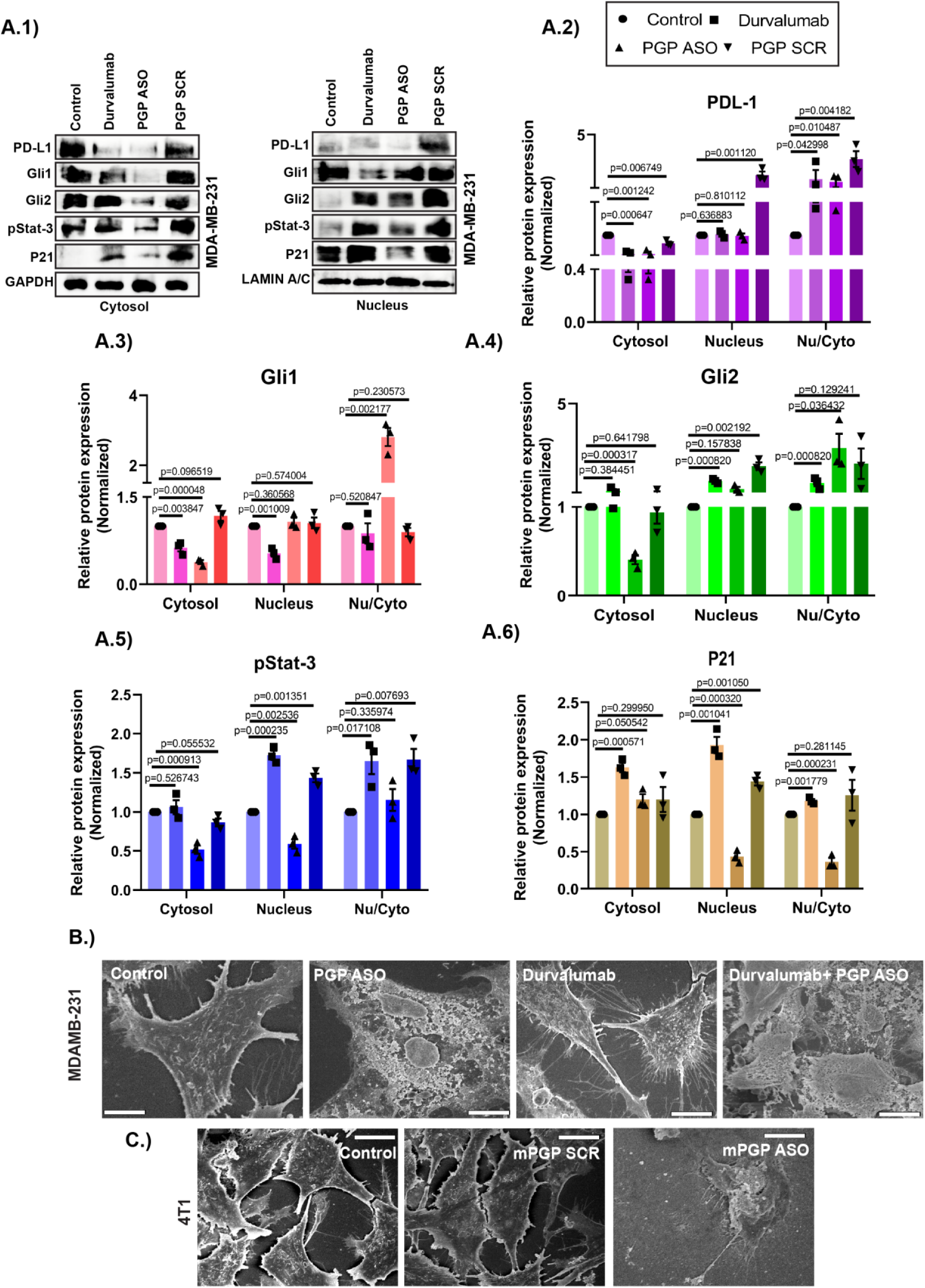
PD-L1 nuclear transportation and ultrastructure modulation compared to durvalumab treatment. **(A1)** Nuclear and cytosolic fractionation of MDA-MB-231 cell shows PD-L1 and associated proteins expression patterns. **(A2-6)** Densitometric analysis of PD-L1, Gli1, Gli2, pStat-3 and P21 proteins respectively after cellular fractionation. **(B)** FE-SEM images of MDA-MB-231 cell after different drug treatments. Scale bar=1μm. **(C)** FE-SEM images of 4T1 cell after mPGP ASO treatment. Scale bar=1μm Data shown as mean ± SEM, Multiple t test was performed followed by Holm-sidak multiple comparison test. *p* ≤ 0.5 considered as statistically significant.

PD-L1 interacts with DNA by its c-tail to regulate different target gene expression^33^. To investigate the localized expression pattern of different transcription factors linked with PD-L1, pStat-3, Gli1, Gli2 and P21 proteins were chosen to investigate. Among these only pStat-3 and P21 showed less nuclear to cytosolic protein expression ratio. Gli1 and Gli2 were downregulated in cytosolic fraction but not significantly in nucleus after PGP ASO treatment (**Figure 4A1-6**).

Besides this, to visualize ultra structural changes in morpholino treated cells, we performed FE-SEM which clearly indicated morphological changes after PGP ASO treatment in comparison to durvalumab (**Figure 4B-C**). Cell to cell connections were hampered in case of PGP ASO treatment but durvalumab showed cell connections which indicated superiority of mRNA level PD-L1 targeted treatment.

### Strategic PD-L1 Targeting in TNBC: Concomitant Disruption of PD-1/PD-L1 Binding *in Vitro* and *in Vivo*

PD-1/PD-L1 engagement induces T-cell exhaustion and apoptosis. Morpholino mediated PD-L1 silencing may reverse this immunosuppression. Co-culture of pre-activated CD4⁺ Jurkat cells with MDA-MB-231 revealed marked PD-L1 downregulation—80% with PGP ASO and 90% with dual PGP ASO–durvalumab treatment (**Figure 5A**). IRF-1 expression was concurrently suppressed. Apoptotic profiling showed a 46% increase in Bax and a 385% elevation in cleaved caspase-9 in the PGP ASO group, indicating robust tumoricidal T-cell activation (**Figure 5 B1,2 & S16A**).

**Figure 5:**
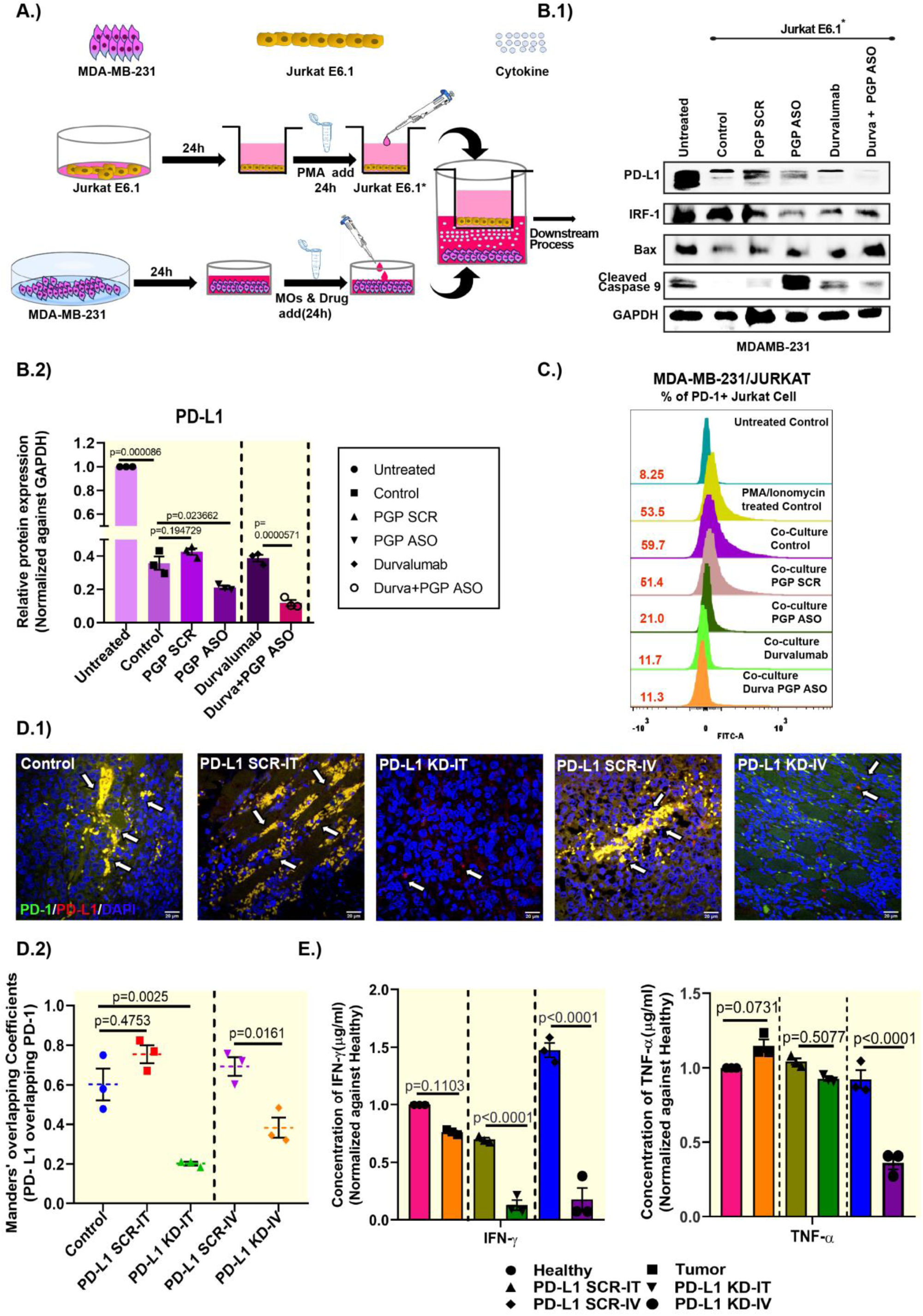
Targeting PD-L1 by PGP ASO alters PD-1 expression *in-vitro* and in-vivo. **(A)** Schematic representation of co-culture assay. **(B.1 & B.2)** Western blot analysis of PD-L1 and associated proteins of MDA-MB-231 cell from coculture system and densitometric analysis of PD-L1 proteins from coculture system. **(C)** Graphical representation of PD-1+ Jurkat cell count after coculture with different treatments on MDA-MB-231 cells. **(D.1 & D.2)** Co-localization of PD-L1 and PD-1 in tumor tissue. Scale bar=20μm.**(E)** Systemic IFN-γ and TNF-α measurement by ELISA in different PD-L1 KD mice. Data shown as mean ± SEM, Multiple t test was performed followed by Holm-sidak multiple comparison test. *p* ≤ 0.5 considered as statistically significant.

PD-1 expression in co-cultured activated Jurkat E6.1* cells was assessed across treatments. In MDA-MB-231, dual PGP ASO–Durvalumab treatment yielded maximal PD-1 suppression (**Figure 5C**). Notably, Jurkat cells received no direct treatment. These results highlight that PD-L1 inhibition indirectly modulates T-cell activity, enhancing antitumor immunity.

Furthermore, in *in vivo* experiments, intertumoral (IT) and intravenous (IV) administration of mPGP ASO revealed that targeting PD-L1 not only downregulates PD-1 but also augments FOXO-3A expression (**Figure S16B**). Additionally, immunofluorescence imaging demonstrated a stark diminution in PD-1/PD-L1 colocalization post-mPGP ASO administration, as illustrated in the corresponding diagram (**Figure 5D1-2**).

### PGP ASO reshape the tumor microenvironment while modulating the immune-cell repertoire

IFN-γ, a key regulator of PD-L1 and marker of immunotherapy response, was reduced following PD-L1 inhibition in PD-L1 KD mice (0.126 ± 0.044 IT; 0.175 ± 0.103 IV) compared to healthy controls, indicating its basal requirement for adaptive immunity. IFN-γ decline reflects its regulatory role in PD-L1/PD-1 signaling. PD-L1 KD-IV mice showed decreased in TNF-α levels by 78% (0.362 ± 0.045), while PD-L1 KD-IT group showed minimal effect (**Figure 5E**). Given TNF-α’s role in T-cell apoptosis and immune evasion, its suppression further underscores the immunomodulatory and therapeutic efficacy of PGP morpholino.

Cytokines can trigger apoptosis within tumors. In tumor-derived neoplastic cells, PD-L1 KD mice showed markedly elevated ROS (via DCFDA) and reduced mitochondrial membrane potential by 49% (via JC-1) (**Figure S17 A & B**). Primary ascites cultures treated with mPGP ASO showed G2-M arrest after 48 hrs. (**Figure S17C**). Additionally, BrdU incorporation revealed significant DNA damage induction in PD-L1 KD tumors (**Figure S17D**), confirming antiproliferative activity.

iNOS, an important M1 macrophage marker, was significantly upregulated in both PD-L1 KD-IT and PD-L1 KD-IV tumors (**Figure 6 A1,2**), indicating modulation of TILs via PD-L1 knockdown. Notably, immunosuppressive regulatory cells (CD4⁺CD25⁺FOXP3⁺) were reduced from 7.1% to 1.42%, and immunosuppressive CD8⁺ regulatory (CD8⁺CD25⁺FOXP3⁺) cells dropped from 8.82% to 0.53% post-treatment (**Figure 6B, C**). Additionally, CD4⁺ and CD8⁺ cells showed enhanced profiles (**Figure 6D**), underscoring the immunotherapeutic efficacy of PGP ASO.

**Figure 6:**
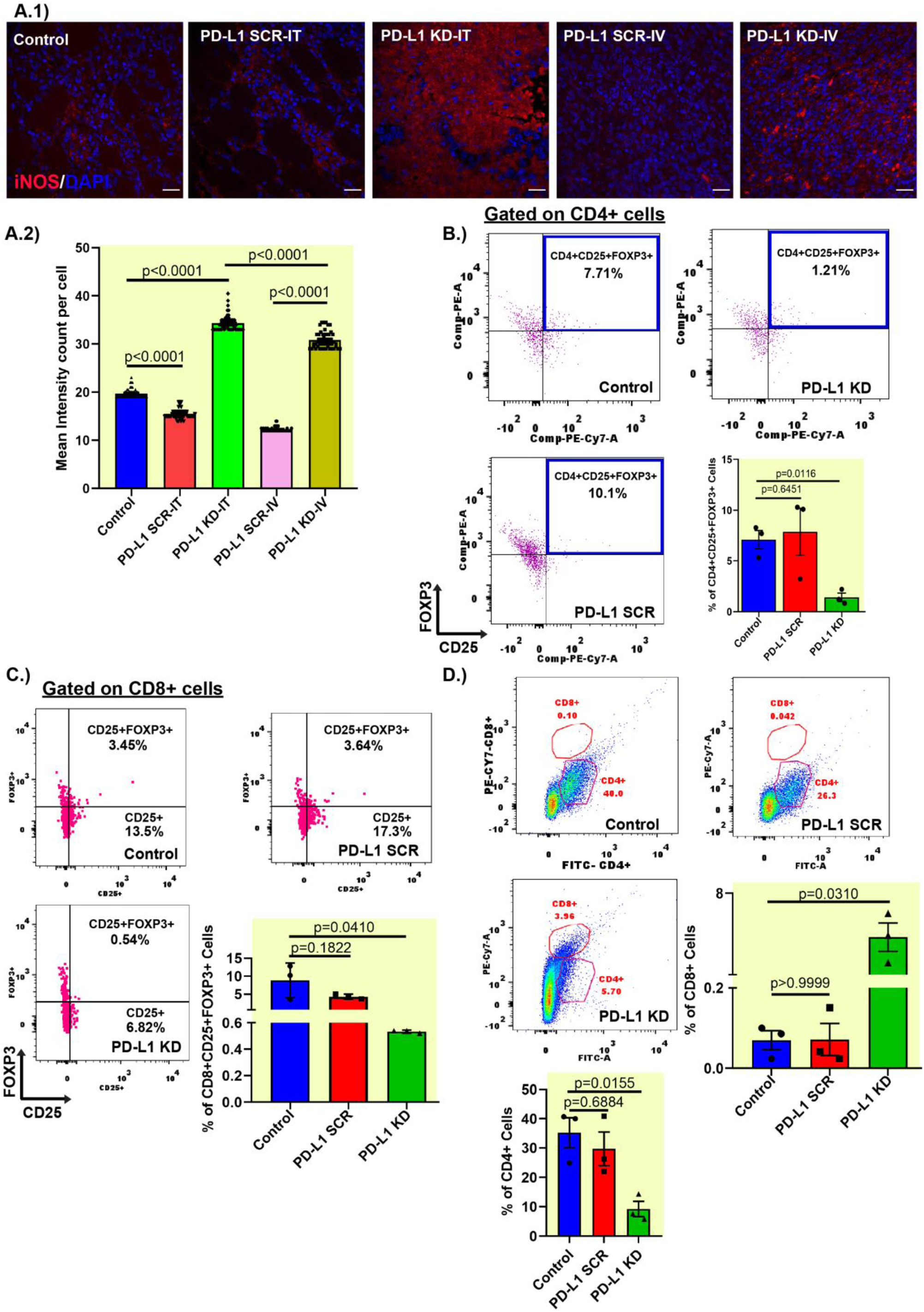
mPGP ASO regulates tumor microenvironment. (**A.1**) iNOS induction after mPGP ASO administration intratumorally and intravenous. (**A.2**) Graphical representation of mean intensity counts of iNOS per cell. Data shown as mean ± SEM, Ordinary one-way ANOVA was performed followed by sidak multiple comparison test. *p* ≤ 0.5 considered as statistically significant. **(B)** Depiction of CD4+CD25+FOXP3+ cell population in tumor of PD-L1 KD/SCR mice. Data shown as means ± SEM, paired parametric t-test was performed. *p* ≤ 0.5 considered as statistically significant. **(C)** Depiction of CD8+CD25+FOXP3+ cell population in tumor of PD-L1 KD/SCR mice. Data shown as means ± SEM, unpaired parametric t-test was performed. *p* ≤ 0.5 considered as statistically significant. **(D)** Flow cytometric analysis of CD4+ and CD8+ cells in whole tumor mass of PD-L1 KD/SCR mice. Ordinary one-way ANOVA was performed followed by sidak multiple comparison test. *p* ≤ 0.5 considered as statistically significant.

### Comprehensive assessment of off-target effects of PGP ASO *in vitro* and *in vivo* models

Off-target effects remain a major challenge in nucleic acid-based therapeutics. Unlike Vivo-morpholino, which induces myocardial toxicity, our GMO-PMO chimera exhibits serum stability and organ safety^34^. mPGP ASO targeting PDL-1 showed no systemic toxicity, with normal CKMB, creatinine, SGPT, SGOT, and albumin levels (**Figure S18 A–F**), and unaltered histopathology, particularly in cardiac tissue.

TLRs, including endosomal TLR7 and TLR9, are expressed in tumor cells and detect exogenous nucleic acids^35, 36^. qRT-PCR analysis showed no significant TLR7 changes, except in PGP ASO-treated MDA-MB-468 (increased 192.67 ± 94.26%) and siRNA-treated MDA-MB-231 cells (increased 150.35 ± 54.09%) (**Figure S18 G**), likely due to PD-L1 suppression-linked TLR upregulation. TLR9 remained unchanged in MDA-MB-468 but was elevated in siRNA and PGP SCR-treated MDA-MB-231 cells (**Figure S18H**), suggesting subtype and treatment-specific modulation.

Caspase activation marks first level of cytotoxicity assessment in *in-vitro* conditions of most of the oligonucleotide’s research field^37^. Assessment of caspase 9 expression indicated cytotoxicity induced by morpholino, whereas other treatments showed no statistically significant upregulation of the same (**Figure S18I**). Altogether, the results further re-established guanidium-linked morpholino oligonucleotide (PGP) mediated targeted regulation of PD-L1 and its downstream pathways (**Figure S18J**). Additionally, mPGP ASO restored thymic architecture in tumor-bearing mice^38^, reestablishing cortex-medulla delineation without altering histopathological architecture **(Figure S19**), supporting its translational safety.

## Discussion

Morpholino oligonucleotides constitute one of the most prevalently employed antisense chemistries in the field of developmental biology. Although four morpholino-based therapeutics have received FDA approval for the treatment of Duchenne Muscular Dystrophy (DMD), their clinical translation into oncology has been impeded, primarily due to suboptimal cellular uptake. To address this limitation, we previously developed guanidinium-modified morpholino oligonucleotides (GMO-PMO chimeras), which exhibit markedly enhanced intracellular permeability^27^. Cell permeable morpholino has not been studied yet in immunotherapeutic uses. In the current investigation, we reported both cellular and murine model evidence demonstrating a targeted, dose-responsive silencing of PD-L1 expression via GMO-PMO administration. Intravenous administration of PGP ASO not only showed significant inhibition of target protein but also exhibited no serum toxicity and vital organ damage.

Immunotherapeutic modalities exhibit inherent limitations in overall response rates. ‘Cold’ neoplasms demonstrate heightened resistance to immune checkpoint interventions^39^. Such refractoriness is frequently attributable to aberrant transcription factor modulation, culminating in attenuated responsiveness to conventional therapeutic regimens.

PD-L1-targeting morpholino PGP induced dose-dependent PD-L1 suppression in two TNBC cell lines, concomitant with significant tumor volume reduction via both intra-tumoral (PD-L1 KD-IT) and systemic delivery (PDL-1 KD-IV). PGP ASO not only suppressed PD-L1 but also disrupted PI3K/Akt and Stat3 signaling, downregulating Cdc42 and p21 while robustly upregulating FOXO-3A, a key NF-κB regulated PD-L1 repressor. Unlike siRNA, which failed to induce FOXO-3A and paradoxically enhanced NF-κB, PGP ASO demonstrated superior specificity and immunomodulatory efficacy, underscoring the therapeutic advantage of morpholino-based antisense platforms. (**Figure 1**).

The cell-penetrating efficiency of morpholino was reaffirmed using PGP ASO, PGP SCR, and PP ASO. Lacking guanidinium, PP ASO failed cellular uptake, whereas PGP ASO and SCR escaped lysosomal compartment within 20 h (**Figure S11**). IFN-γ activates the JAK2-Stat3 axis via receptor phosphorylation, facilitating Stat3 nuclear translocation and IRF-1 mediated PD-L1 transcription^40^. This study recapitulated the IFN-γ pathway to evaluate PGP ASO efficacy. Notably, PGP ASO not only suppressed downstream PD-L1 signaling but also induced autophagic cell death. In MDA-MB-231, LC3B was upregulated and Beclin downregulated; MDA-MB-468 exhibited inverse expression, indicating model-dependent functionality, corroborated by *in vivo* data (**Figure 2**).

N-glycosylation governs PD-L1 surface trafficking^41^ via endocytosis, macropinocytosis, and lipid raft-mediated routes. Genistein (in MDA-MB-231) and Mβ-CD (in MDA-MB-468) most effectively disrupted PD-L1 recycling. Confocal analysis showed that PGP ASO suppressed PD-L1 surface expression comparably to durvalumab while achieving superior mRNA knockdown, with downstream effectors following expected inhibitory trends (**Figure 3**).

Nuclear PD-L1 sustains its feedback loop and drives immunotherapy resistance. PGP ASO reduced cytosolic Gli1, Gli2, pStat3, and p21, while residual nuclear Gli1/2 indicated limited morpholino nuclear entry. Nuclear PD-L1 persisted only in the PGP SCR group, whereas durvalumab suppressed Gli1 but spared other transcription factors. Additionally, ultrastructural aberrations within malignant cells were evident post-PGP ASO treatment, both as monotherapy and in combination with durvalumab (**Figure 4**).

PGP ASO mediated PD-L1 suppression enhanced T cell mediated cytotoxicity *in vitro* and *in vivo*, upregulating pro-apoptotic Bax and cleaved caspase-9. Co-culture with Jurkat E6.1 cells showed maximal PD-1 attenuation in MDA-MB-231 cells treated with PGP ASO plus durvalumab, overcoming T cell exhaustion. *In vivo*, systemic mPGP ASO markedly reduced PD-L1/PD-1 co-localization (Manders overlapping coefficient), confirming both PD-L1 inhibition and disruption of PD-L1/PD-1 binding (**Figure 5**).

IFN-γ and TNF-α serve as critical biomarkers of tumor burden. Notably, TNF-α confers immunosuppressive resistance by promoting apoptosis of cytotoxic T lymphocytes^42^, an effect substantially abrogated following intravenous administration of mPGP ASO (**Figure 5B**). Moreover, inducible nitric oxide synthase (iNOS), a hallmark of M1-polarized macrophages^43^, exhibited pronounced upregulation following intra-tumoral delivery of mPGP ASO. The immunosuppressive tumor microenvironment, predominantly orchestrated by Treg cells, facilitates immune evasion via granzyme B and perforin mediated lysis of cytotoxic T lymphocytes. CD4⁺CD25⁺FOXP3⁺ (or CD8^+^CD25⁺FOXP3⁺) effector regulatory cells exert robust immunomodulatory control through both adaptive and humoral axes, secreting immunosuppressive cytokines such as IL-10 and TGF-β^44, 45^. Flow cytometric analysis of CD4⁺, CD8⁺, and CD25⁺FOXP3⁺ (CD4+ or CD8+) populations revealed a marked reduction in regulatory cells abundance in PDL-1 KD mice (**Figure 6**). This depletion likely reflects mPGP ASO induced intra-tumoral reactive oxygen species (ROS) accumulation and DNA damage, collectively contributing to selective immunosuppressive regulatory cell ablation within the tumor niche (**Figure S17**).

Oligonucleotide-based therapeutics are frequently associated with off-target hybridization, potentially eliciting systemic toxicity and consequent damage to vital organs. Commercially available formulations such as Vivo-morpholino have been reported to induce cardiotoxicity, impairing myocardial integrity. In this study, we assessed serum biochemical parameters in mPGP ASO–treated mice (PD-L1 KD-IT and PD-L1 KD-IV) relative to healthy controls. No significant deviations were observed following the treatment regimen. Furthermore, histopathological examination of critical organs, including cardiac tissue, revealed no evidence of chronic toxicity post mPGP ASO administration (**Figure S18 & S19**). Complementary analyses, including TLR7/9 activation and Caspase-Glo 9 assays, further corroborate the systemic safety of guanidinium-linked morpholino (PGP) administration.

Despite these promising results, our study has several limitations. We administered mPGP ASO intravenously, which does not enable selective tumor uptake. The tumor-selective biodistribution profile of mPGP ASO remains under investigation and will be addressed in future studies. Moreover, PGP ASO exhibits limited nuclear transfection, potentially restricting transcription factor modulation.

To enhance cellular uptake, Gene Tools developed Vivo-PMO^46^ by appending arginine-like flexible guanidinium groups to the PMO, enabling sequence-specific gene silencing in cancer models^47^. In contrast, our GMOPMO chimera integrates guanidinium linkages directly into the backbone in a spatially ordered manner, minimizing non-specific interactions and circumventing ASO–associated toxicities^48^. Furthermore, solubility of the chimera into water or PBS, makes it more favorable in use while its 3′ terminus can be readily modified to confer cancer-cell selectivity. Lastly its easy synthesis, favorable cell permeability, cytosolic mRNA target specificity, and serum stability open new avenues for future immunotherapy research.

## Data availability

Data will be made available on request.

## Acknowledgments

S.S. thanks SERB SUPRA SPR/2023/000358 and Indian Association for the Cultivation of Science for financial support. S. Sarkar and U.G thank CSIR for their fellowships, S.P thanks IACS for their fellowships.

## CRediT authorship contribution statement

Conceptualization: SSinha, Methodology: SSarkar, UG, SP, Investigation: SSarkar, UG, Visualization: SSarkar, UG, Supervision: SSinha, Writing—original draft: SSinha, SSarkar, Writing—review & editing: SSinha, SSarkar, UG

## Supplementary Information

**Figure S1:**
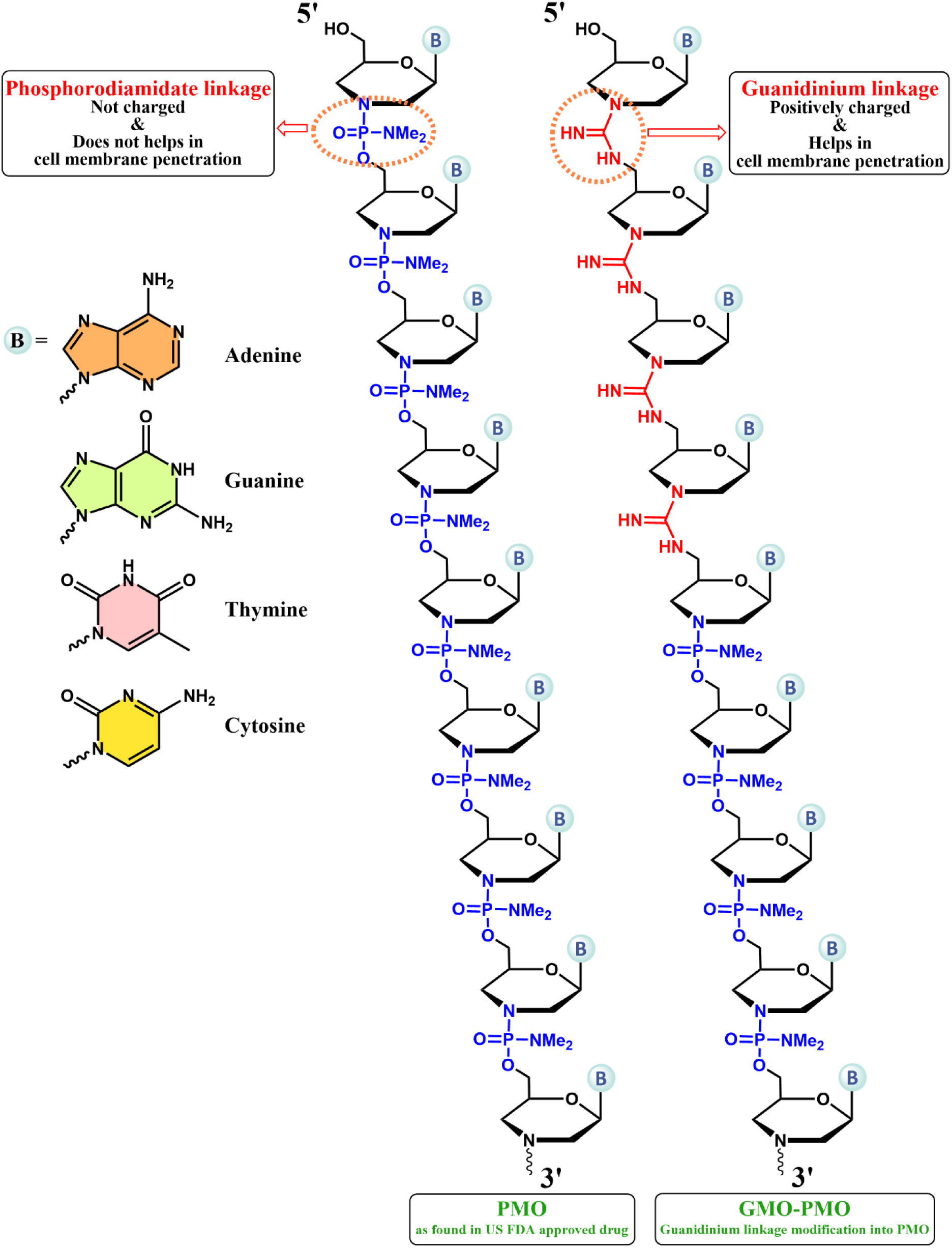
Chemical structure representing. (A) the regular PMO oligomers along with its Phosphorodiamidate backbone and (B) the four guanidinium linkages containing PMO oligomers which are generally termed as GMO-PMO oligomers (Chimera) and are used for this study. Figure S1 is aimed to give a clear picture about how GMO-PMOs are different from PMOs.

**Figure S2:**
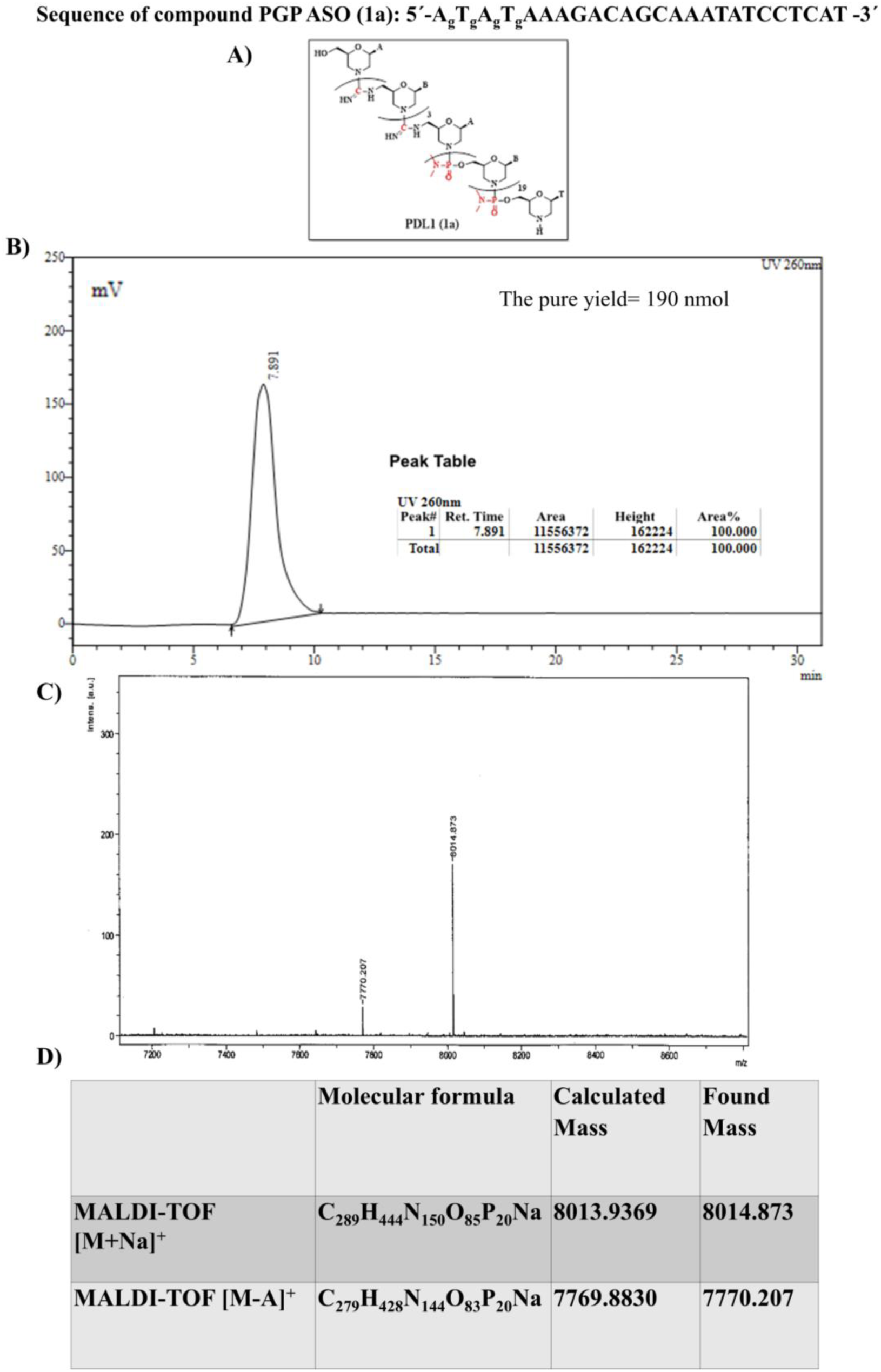
Structure and characterization of PGP ASO. A) Chem-draw structure of PGP ASO. B) HPLC analysis of Pure Compound 1a: Reverse phase C18 column (Xbridge RP18, 10 mm I.D x 250 mm); 20‒50 % CH_3_CN in 0.1 M Ammonium acetate (pH = 7.12) for 20 min and then 50‒20 % CH_3_CN in 0.1 M ammonium acetate for 10 min, flow rate 2 ml/min, R_t_ = 7.891 min. C&D) MALDI-TOF mass analysis of PGP ASO

**Figure S3:**
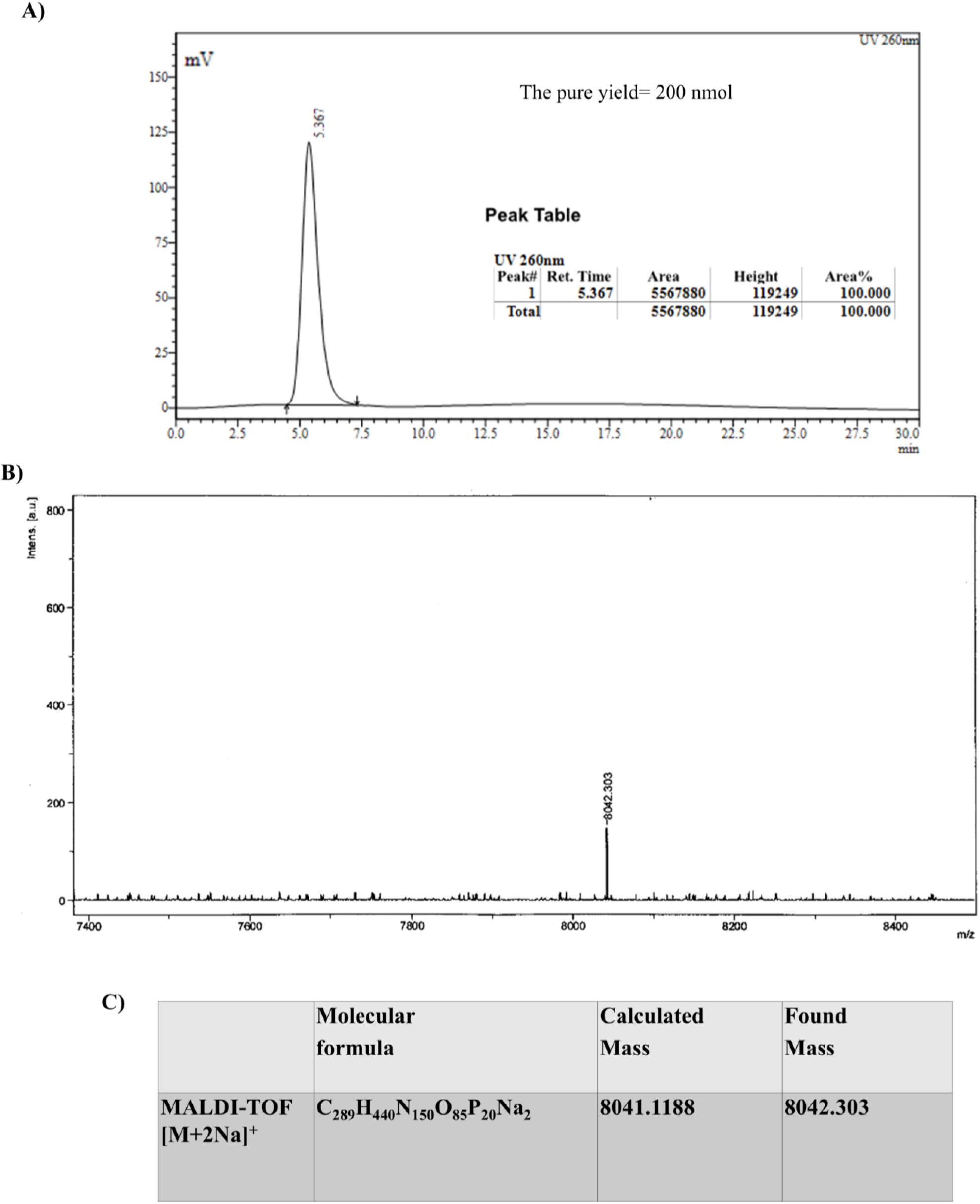
Characterization of PGP SCR. A) HPLC analysis of Pure PGP SCR: Reverse phase C18 column (Xbridge RP18, 10 mm I.D x 250 mm); 20‒50 % CH_3_CN in 0.1 M Ammonium acetate (pH = 7.12) for 20 min and then 50‒20 % CH_3_CN in 0.1 M ammonium acetate for 10 min, flow rate 2 ml/min, R_t_ = 5.367 min. B&C) MALDI-TOF mass analysis of PGP SCR

**Figure S4:**
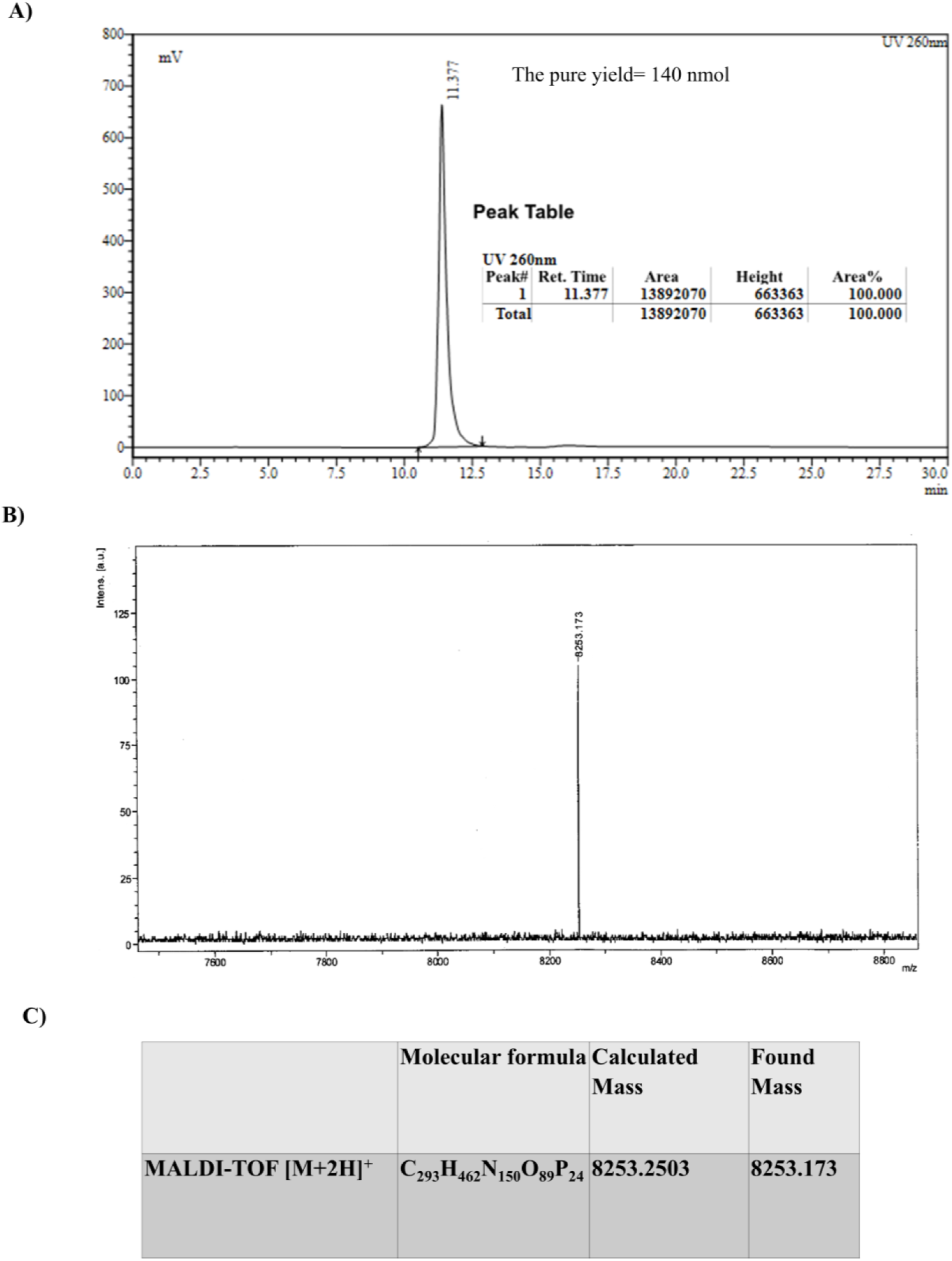
Characterization of PP ASO. A) HPLC analysis of Pure PP ASO: Reverse phase C18 column (Xbridge RP18, 10 mm I.D x 250 mm); 20‒50 % CH_3_CN in 0.1 M Ammonium acetate (pH = 7.12) for 20 min and then 50‒20 % CH_3_CN in 0.1 M ammonium acetate for 10 min, flow rate 2 ml/min, R_t_ = 11.377 min. B&C) MALDI-TOF mass analysis of PP ASO

**Figure S5:**
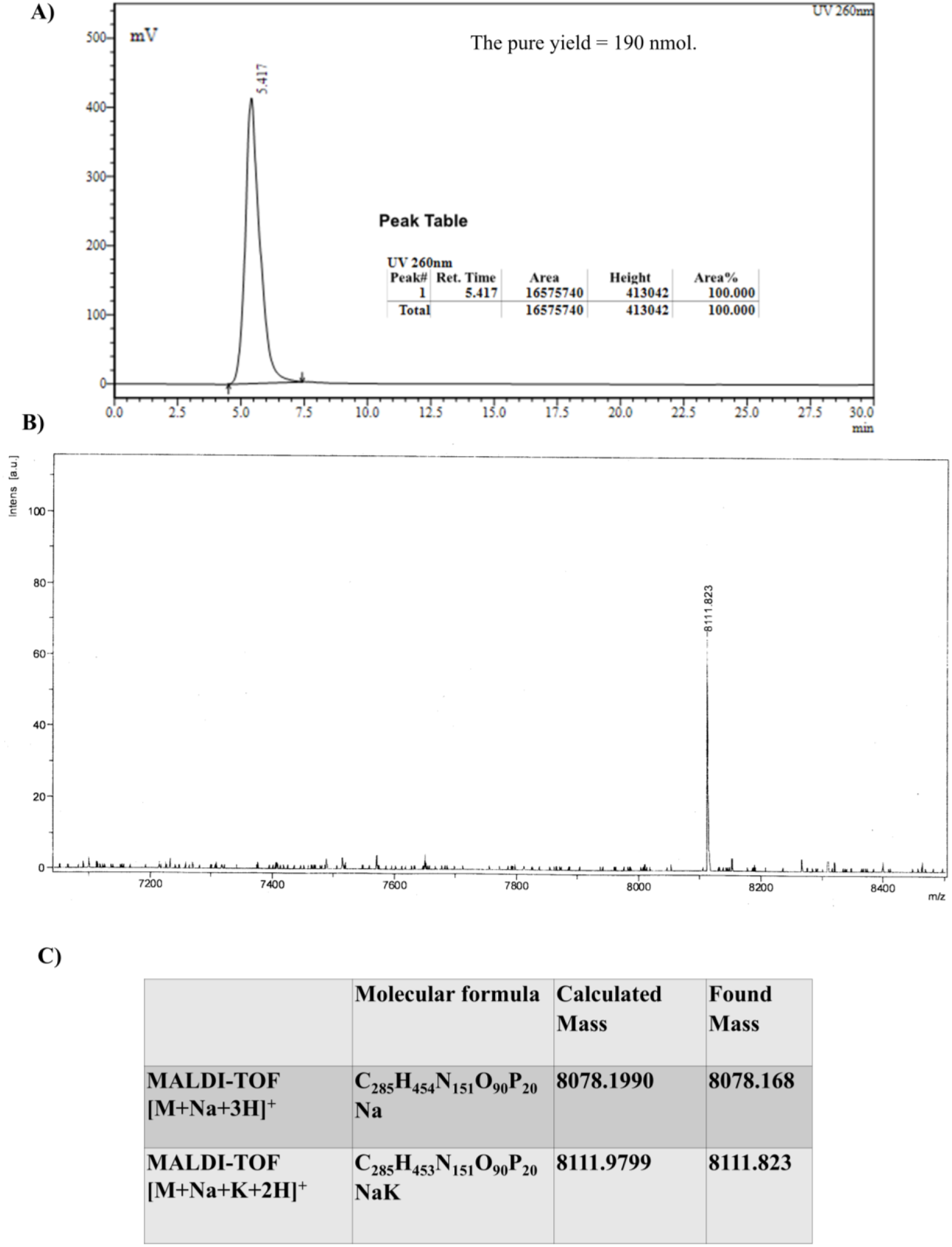
Characterization of mPGP ASO. A) HPLC analysis of Pure mPGP ASO: Reverse phase C18 column (Xbridge RP18, 10 mm I.D x 250 mm); 20‒50 % CH_3_CN in 0.1 M Ammonium acetate (pH = 7.12) for 20 min and then 50‒20 % CH_3_CN in 0.1 M ammonium acetate for 10 min, flow rate 2 ml/min, R_t_ = 5.417 min. B&C) MALDI-TOF mass analysis of mPGP ASO

**Figure S6:**
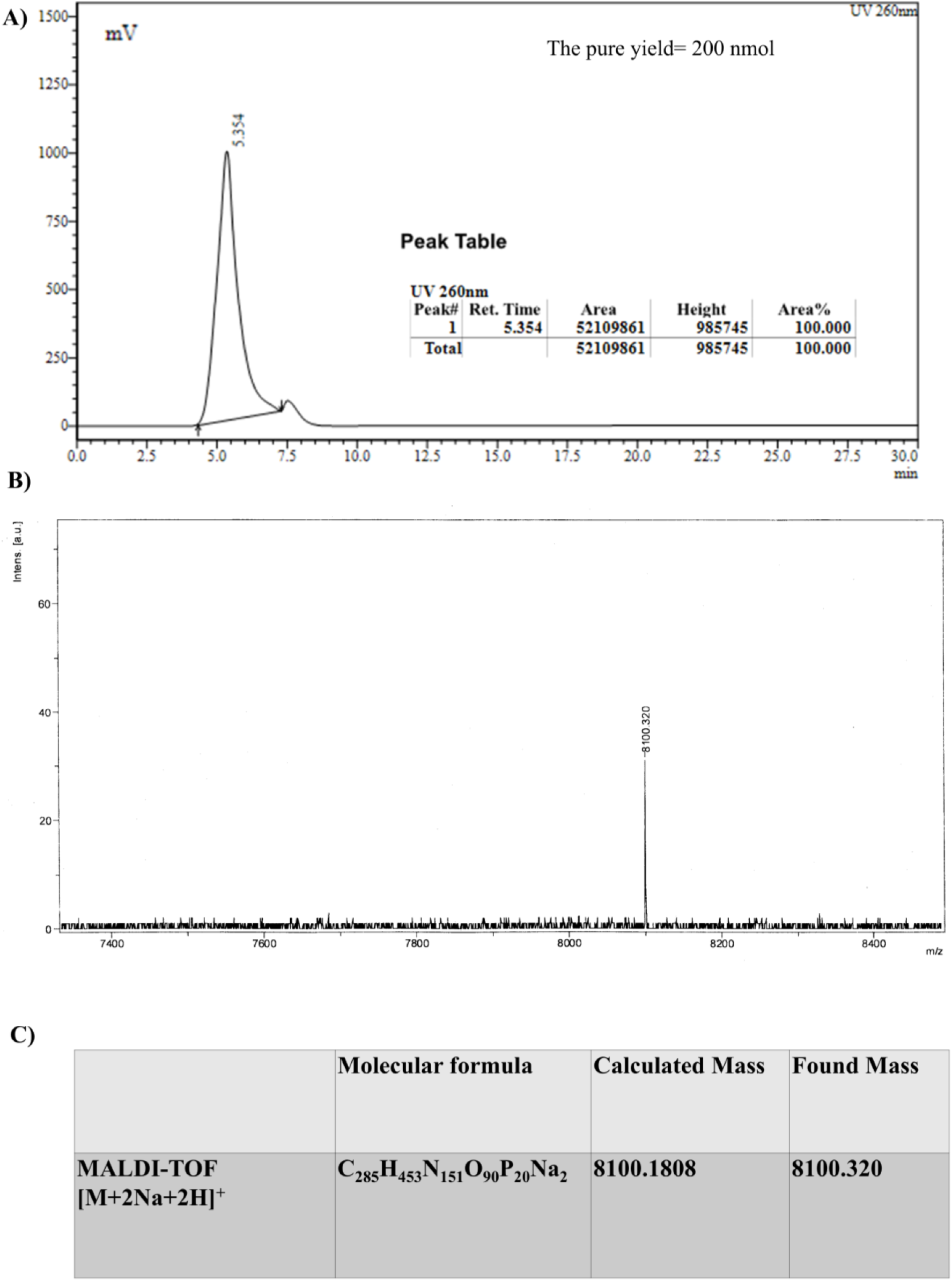
Characterization of mPGP SCR. A) Reverse phase C18 column (Xbridge RP18, 10 mm I.D x 250 mm); 20‒50 % CH_3_CN in 0.1 M Ammonium acetate (pH = 7.12) for 20 min and then 50‒20 % CH_3_CN in 0.1 M ammonium acetate for 10 min, flow rate 2 ml/min, R_t_ = 5.354 min. B&C) MALDI-TOF mass analysis of mPGP SCR.

**Figure S7:**
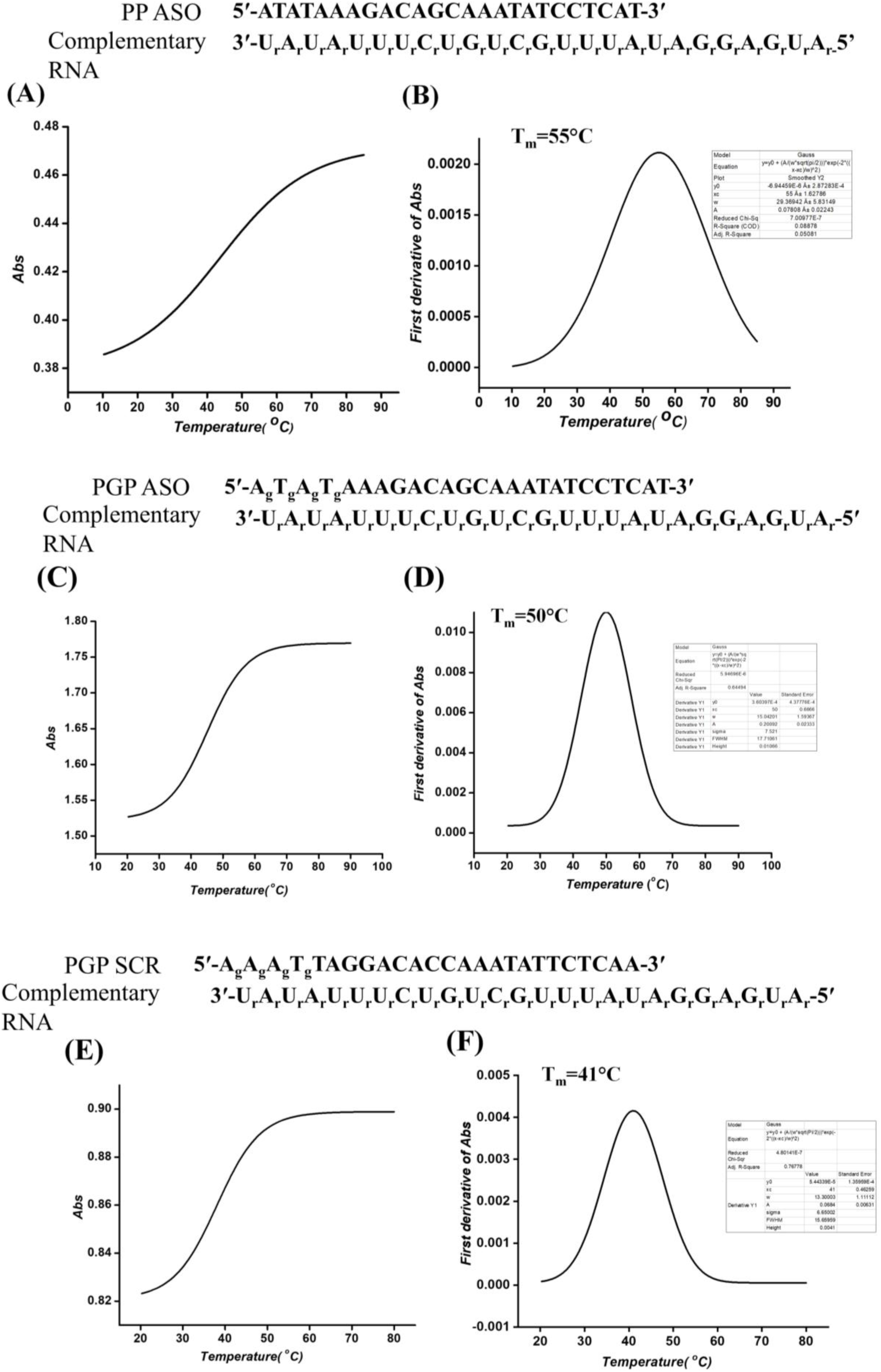
Thermal Melting Curves of morpholino oligonucleotides with complementary RNA. A) Absorption vs. temperature plot and (B) First derivative of absorption vs. temperature plot of Thermal melting in 40 mM phosphate buffer (pH 7) of PP ASO with complimentary RNA. (C) Absorption vs. temperature plot and (D) First derivative of absorption vs. temperature plot of Thermal melting in 40 mM phosphate buffer (pH 7) of PGP ASO with complimentary RNA. (E) Absorption vs. temperature plot and (F) First derivative of absorption vs. temperature plot of Thermal melting in 40 mM phosphate buffer (pH 7) of PGP SCR with complimentary RNA.

**Figure S8:**
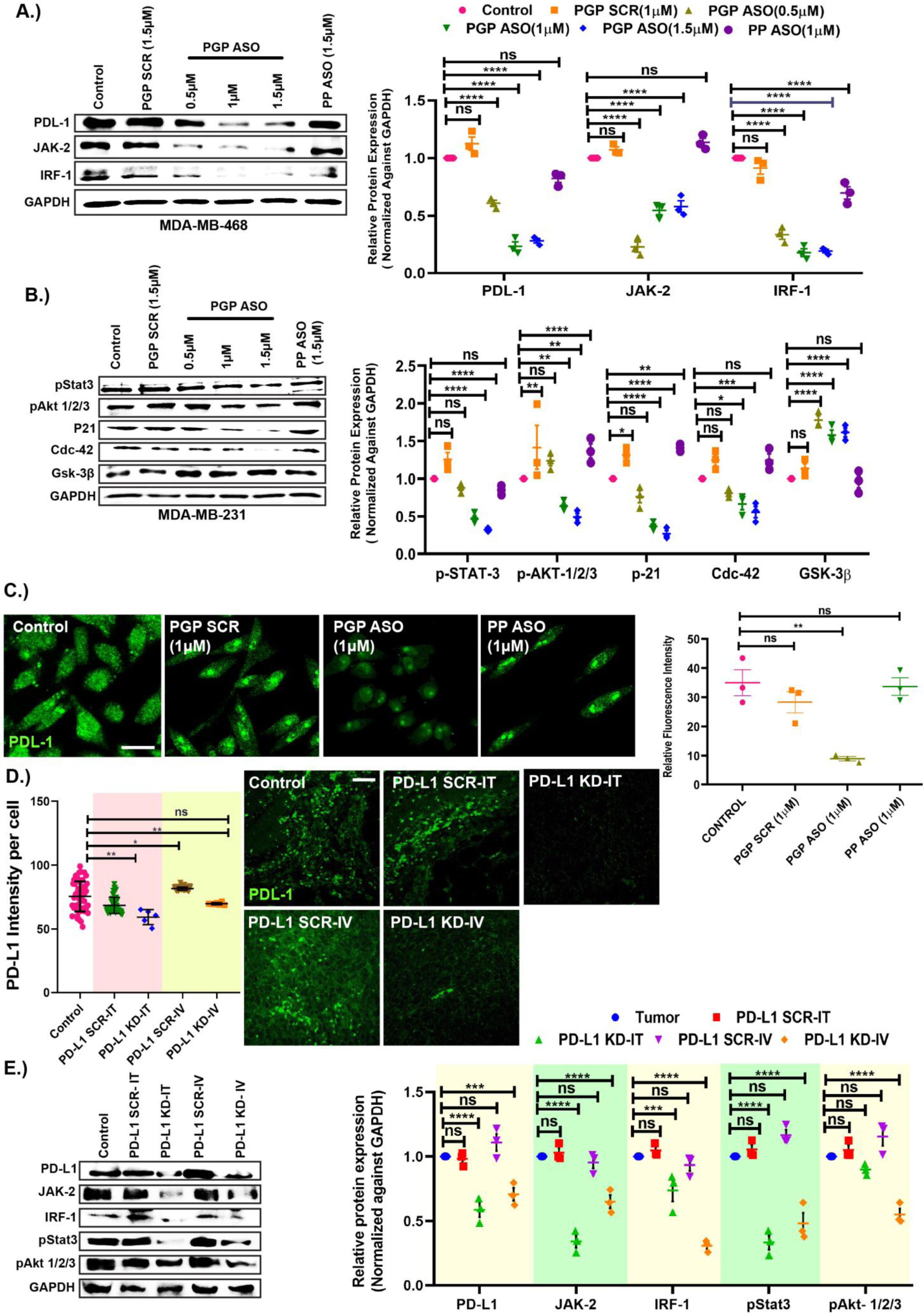
PD-L1 targeted inhibition by cell permeable GMOPMO *in vitro* and *in vivo*. **(A)** Dose dependent inhibition of PD-L1 and associated proteins by PGP ASO in MDA-MB-468 cell (Left panel), Graphical representation of densitometric analysis of PDL-1, JAK-2 and IRF-1 proteins (Right panel). **(B)** Dose dependent inhibition of pStat-3, pAkt 1/2/3, P21, Cdc-42, Gsk-3β in MDA-MB-231 cells. **(C)** Immunocytochemistry of PD-L1 after PGP ASO and PP ASO treatment. Scale bar = 10μM. And statistical representation of relative fluorescence intensity of PD-L1. **(D)** Statistical representation of relative fluorescence intensity of PD-L1 in tumour tissue and immunofluorescence image of PD-L1 in tumour tissue after intra-tumoral and intravenous administration of mPGP ASO. Scale bar = 20μM. **(E)** western blot analysis of PD-L1 and associated protein after mPGP ASO administration (left panel). And graphical representation of densitometric analysis of above said proteins (right panel). Data shown as mean ± SEM, Ordinary Two-way ANOVA was performed followed by sidak multiple comparison test. \**p* ≤ 0.5 considered as statistically significant, ** and *** considered as higher significant than *p* ≤ 0.5.

**Figure S9:**
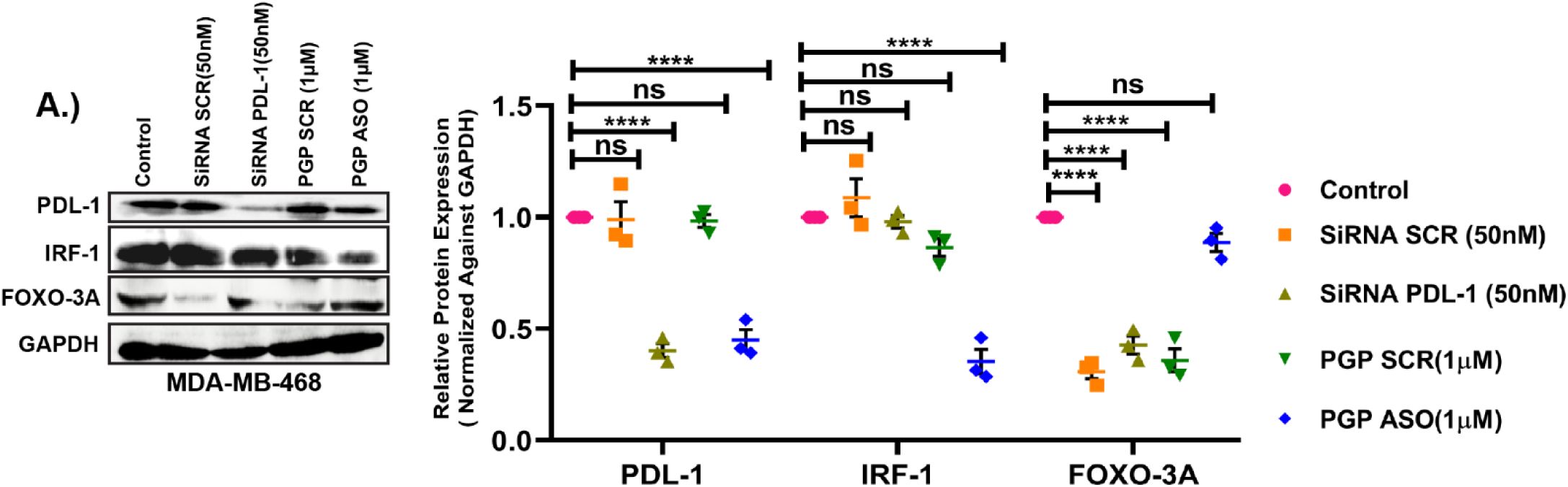
Comparative study with siRNA in MDA-MB-468 cells. **(A)**Representative images of western blot analysis of PGP ASO and siRNA mediated comparative inhibition of PD-L1 and associated proteins in MDA-MB-468(left panel). And Bar diagram representation of densitometric analysis of PGP ASO and siRNA mediated comparative inhibition profiles (right panel). Data shown as mean ± SEM, Ordinary Two-way ANOVA was performed followed by sidak multiple comparison test. \**p* ≤ 0.5 considered as statistically significant, ** and *** considered as higher significant than *p* ≤ 0.5.

**Figure S10:**
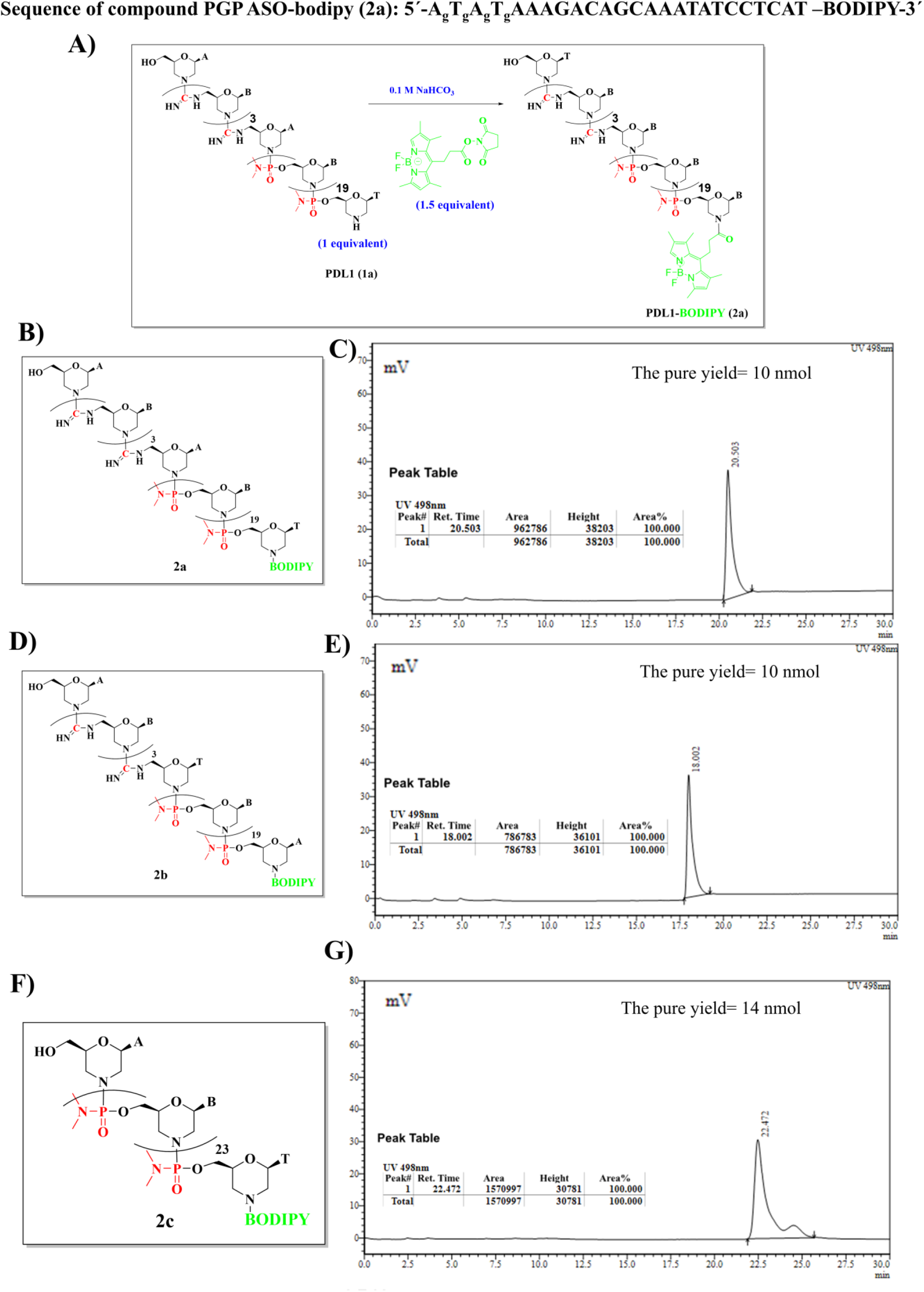
Conjugation of BODIPY with morpholino oligonucleotides. **(A)**Synthesis scheme of bodipy conjugation **(B &C)** Structure and HPLC analysis of PGP ASO-bodipy. Reverse phase C18 column (Xbridge RP18, 10 mm I.D x 250 mm); 20‒50 % CH_3_CN in 0.1 M Ammonium acetate (pH = 7.12) for 20 min and then 50‒20 % CH_3_CN in 0.1 M ammonium acetate for 10 min, flow rate 2 ml/min, R_t_ = 20.53 min. **(D&E)** Structure and HPLC analysis of PGP SCR-bodipy. Reverse phase C18 column (Xbridge RP18, 10 mm I.D x 250 mm); 20‒50 % CH_3_CN in 0.1 M Ammonium acetate (pH = 7.12) for 20 min and then 50‒20 % CH_3_CN in 0.1 M ammonium acetate for 10 min, flow rate 2 ml/min, R_t_ = 18.002 min. **(F&G)** Structure and HPLC analysis of PGP SCR-bodipy. Reverse phase C18 column (Xbridge RP18, 10 mm I.D x 250 mm); 20‒50 % CH_3_CN in 0.1 M Ammonium acetate (pH = 7.12) for 20 min and then 50‒20 % CH_3_CN in 0.1 M ammonium acetate for 10 min, flow rate 2 ml/min, R_t_ = 22.472 min.

**Figure S11:**
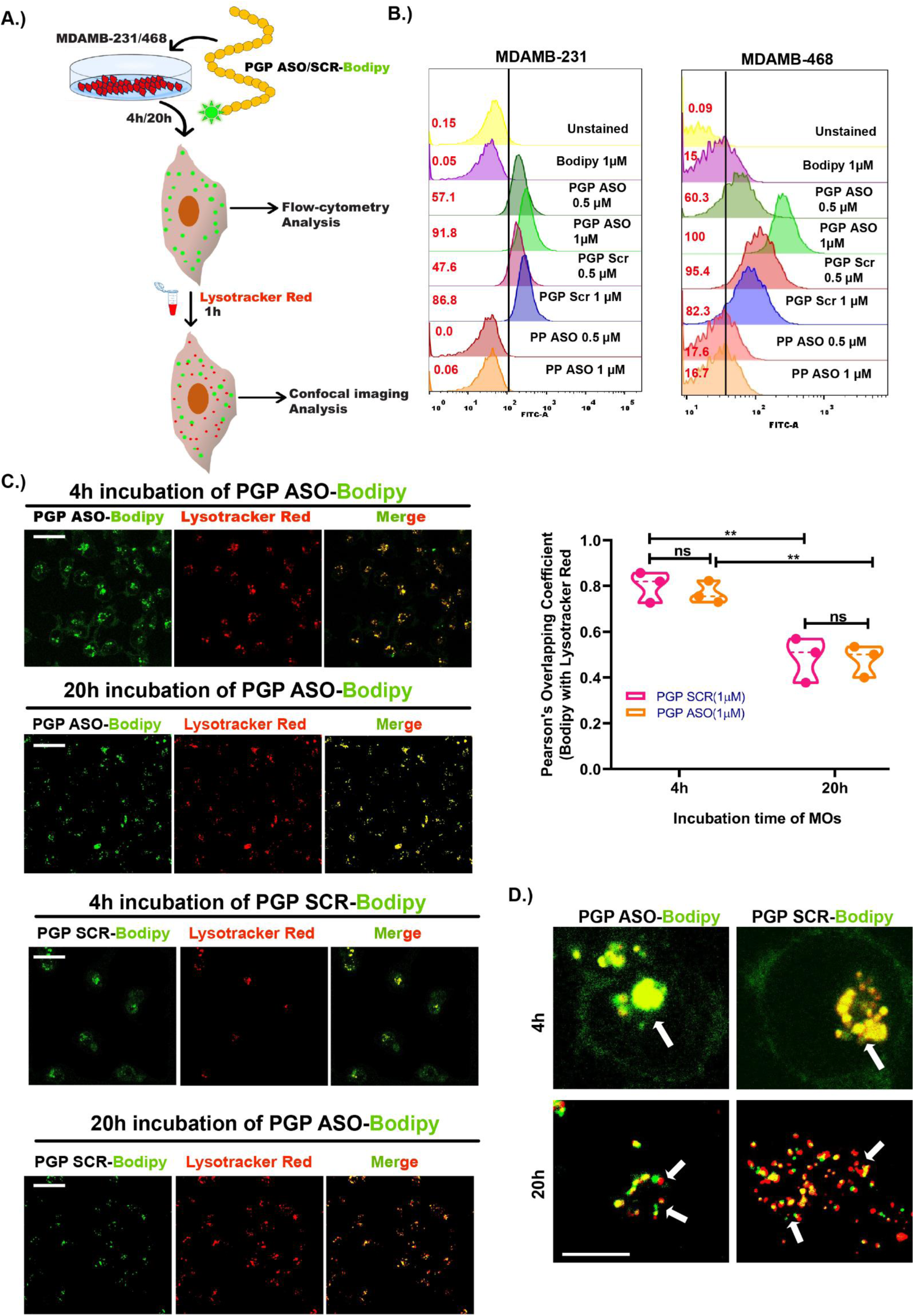
PGP ASO escape the lysosome to enhance PD-L1 inhibition. **(A)** Illustration of endosomal escape and transfection of PGP ASO study details in material and methods section. **(B)** Transfection of PGP ASO-bodipy in MDA-MB-231 and MDA-MB-468 cell lines in comparison with unconjugated bodipy, PGP SCR and PP ASO. **(C)** Confocal images of PGP ASO and PGP SCR distribution in cells in two time points (4h & 20h) and statistical representation of Pearsons’ overlapping coefficient Scale bar=30μm **(D)** Confocal images show (White arrows) PGP ASO/PGP SCR-bodipy escape from lysosome marked by lysotracker red after 20h incubation. Scale bar=10μm Data shown as mean ± SEM, Ordinary Two-way ANOVA was performed followed by sidak multiple comparison test. \**p* ≤ 0.5 considered as statistically significant, ** and *** considered as higher significant than *p* ≤ 0.5.

**Figure S12:**
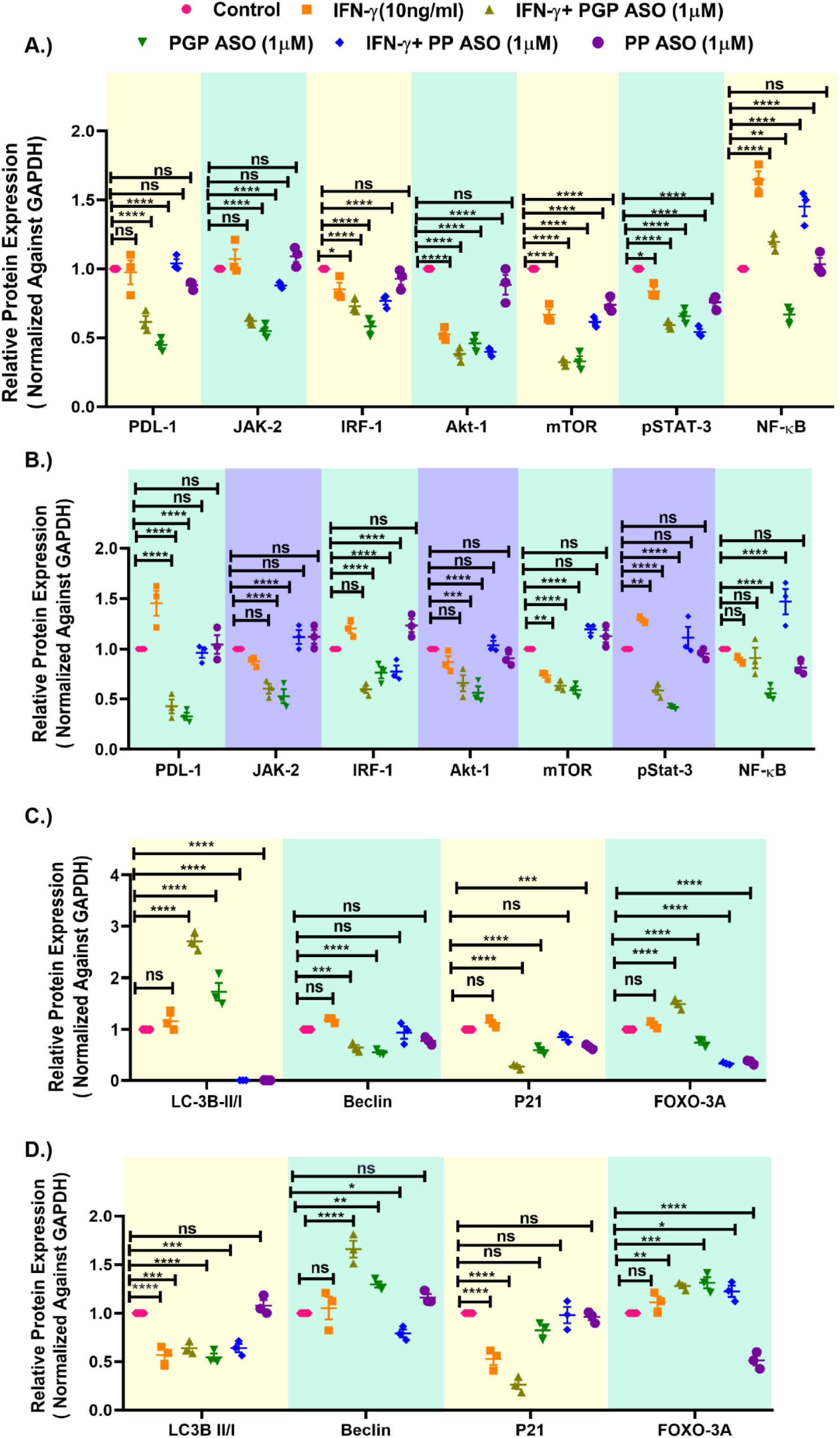
IFN-γ mediated induction of PD-L1 in TNBC. (**A**) Graphical representation of densitometric analysis of figure 2A. **(B)** Graphical representation of densitometric analysis of figure 2B. **(C)** Graphical representation of densitometric analysis of figure 2C. **(D)** Graphical representation of densitometric analysis of figure 2D. Data shown as mean ± SEM, Ordinary Two-way ANOVA was performed followed by sidak multiple comparison test. \**p* ≤ 0.5 considered as statistically significant, ** and *** considered as higher significant than *p* ≤ 0.5.

**Figure S13:**
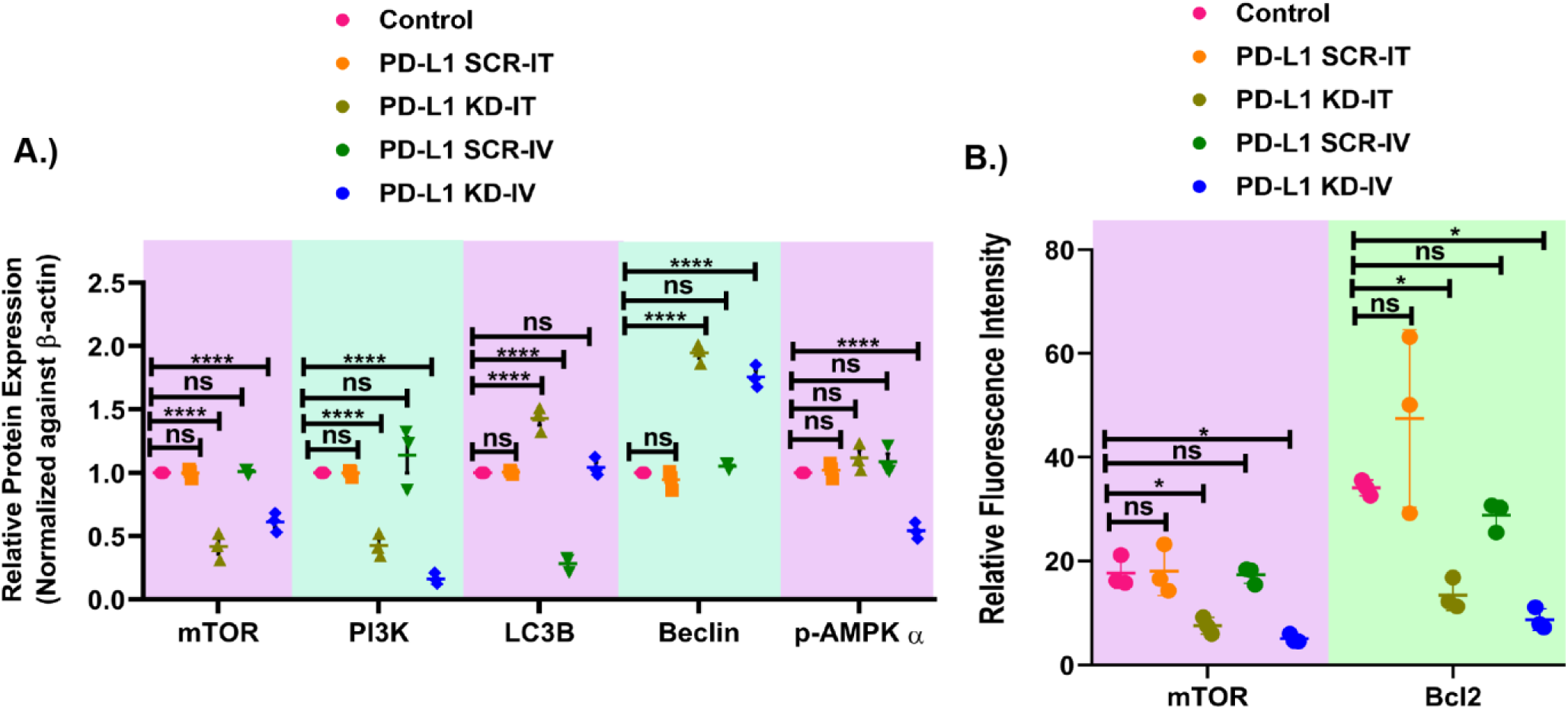
Autophagy pathway assessment *in-vivo*. **(A)** Graphical representation of densitometric analysis of figure 2E. **(B)** Relative fluorescence Intensity of mTOR and Bcl2 of figure 2F image. Data shown as mean ± SEM, Ordinary Two-way ANOVA was performed followed by sidak multiple comparison test. \**p* ≤ 0.5 considered as statistically significant, ** and *** considered as higher significant than *p* ≤ 0.5.

**Figure S14:**
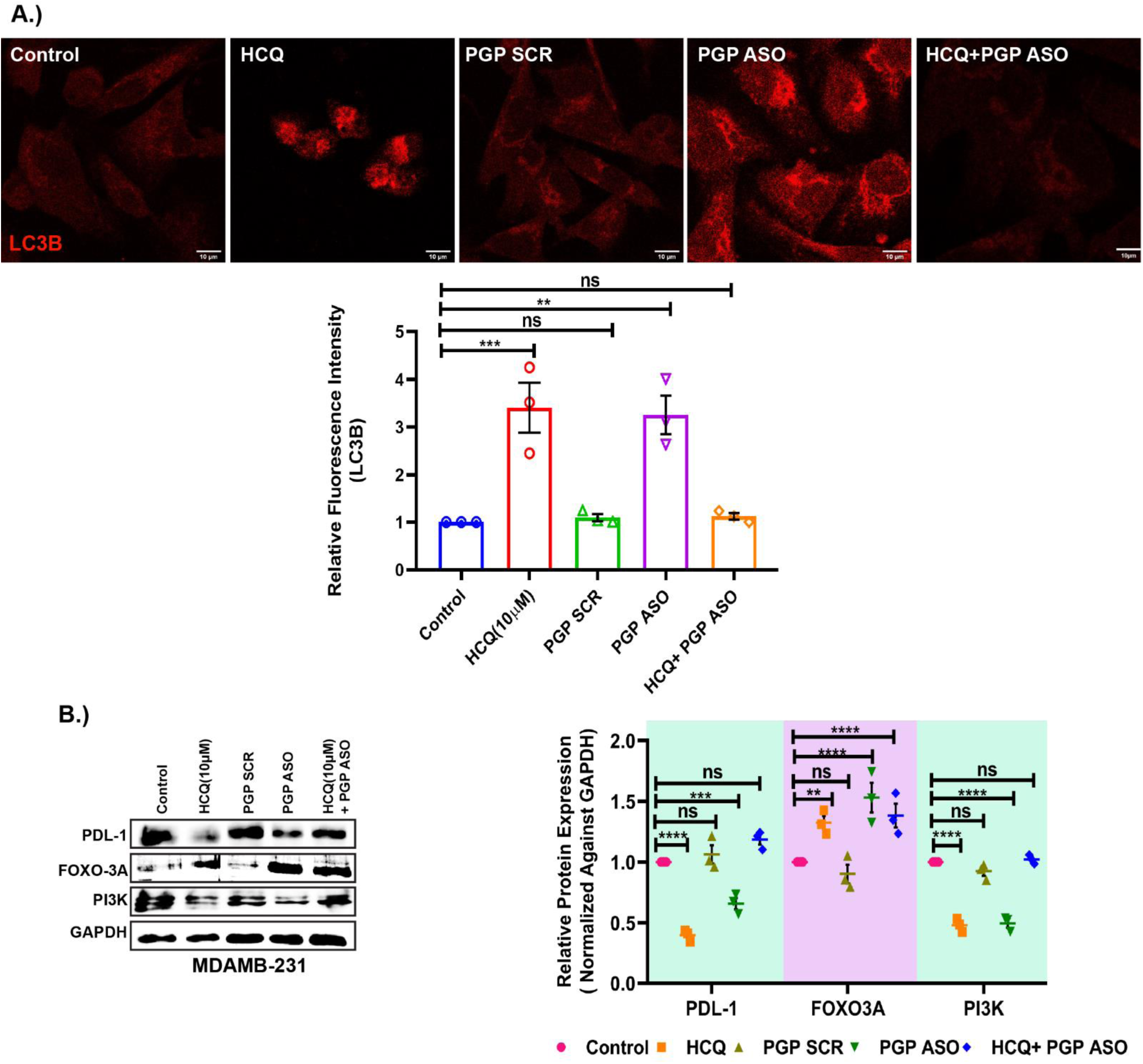
PGP ASO induce autophagy by upregulating LC3B. **(A)**Immunocytochemistry of LC3B after PGP ASO treatment alone and combination with HCQ. Scale bar = 10μm. **(B)** Protein expression profile of PD-L1 and associated proteins in MDA-MB-231 cell with densitometric analysis of above-mentioned proteins. Data shown as mean ± SEM, Ordinary Two-way ANOVA was performed followed by sidak multiple comparison test. \**p* ≤ 0.5 considered as statistically significant, ** and *** considered as higher significant than *p* ≤ 0.5.

**Figure S15:**
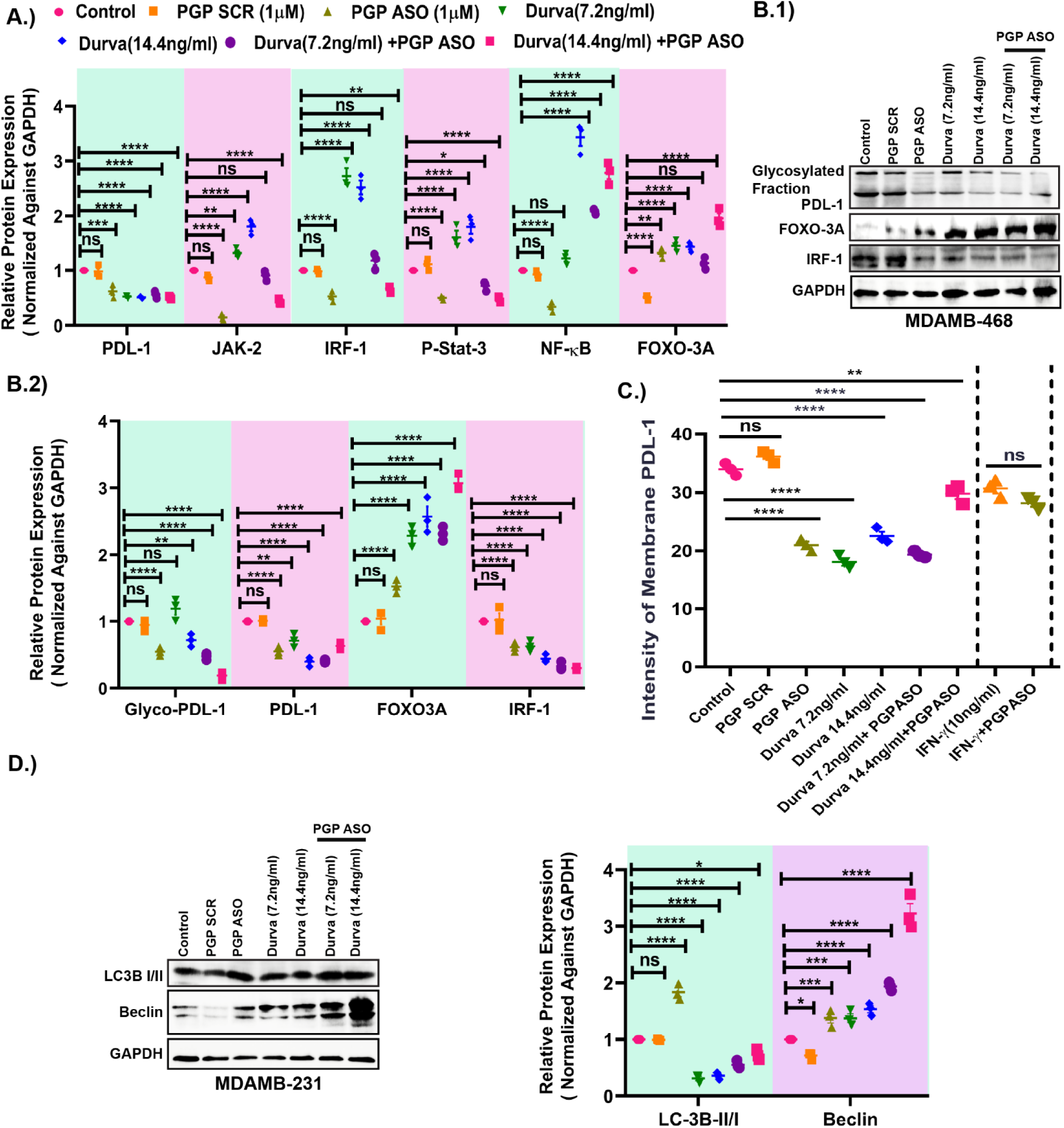
PGP ASO controls glycosylated PD-L1 and associated proteins in combination with durvalumab. **(A)** Graphical representation of densitometric analysis of figure 3B. **(B.1 &B.2)** Western blot analysis of PD-L1 with its glycosylated fraction, FOXO3A and IRF-1 proteins in MDA-MB-468 cells and relative protein expression of glycosylated PD-L1 and associated proteins after PGP ASO treatment alone and in combination with durvalumab. **(C)** Relative fluorescence intensity of cell membrane PD-L1 after different treatment conditions represented in figure 3C. **(D)** Expression pattern of different autophagosomal proteins after PGP ASO and durvalumab treatment in MDA-MB-231 cell and graphical representation of above-mentioned proteins. Data shown as mean ± SEM, Ordinary Two-way ANOVA was performed followed by sidak multiple comparison test. \**p* ≤ 0.5 considered as statistically significant, ** and *** considered as higher significant than *p* ≤ 0.5.

**Figure S16:**
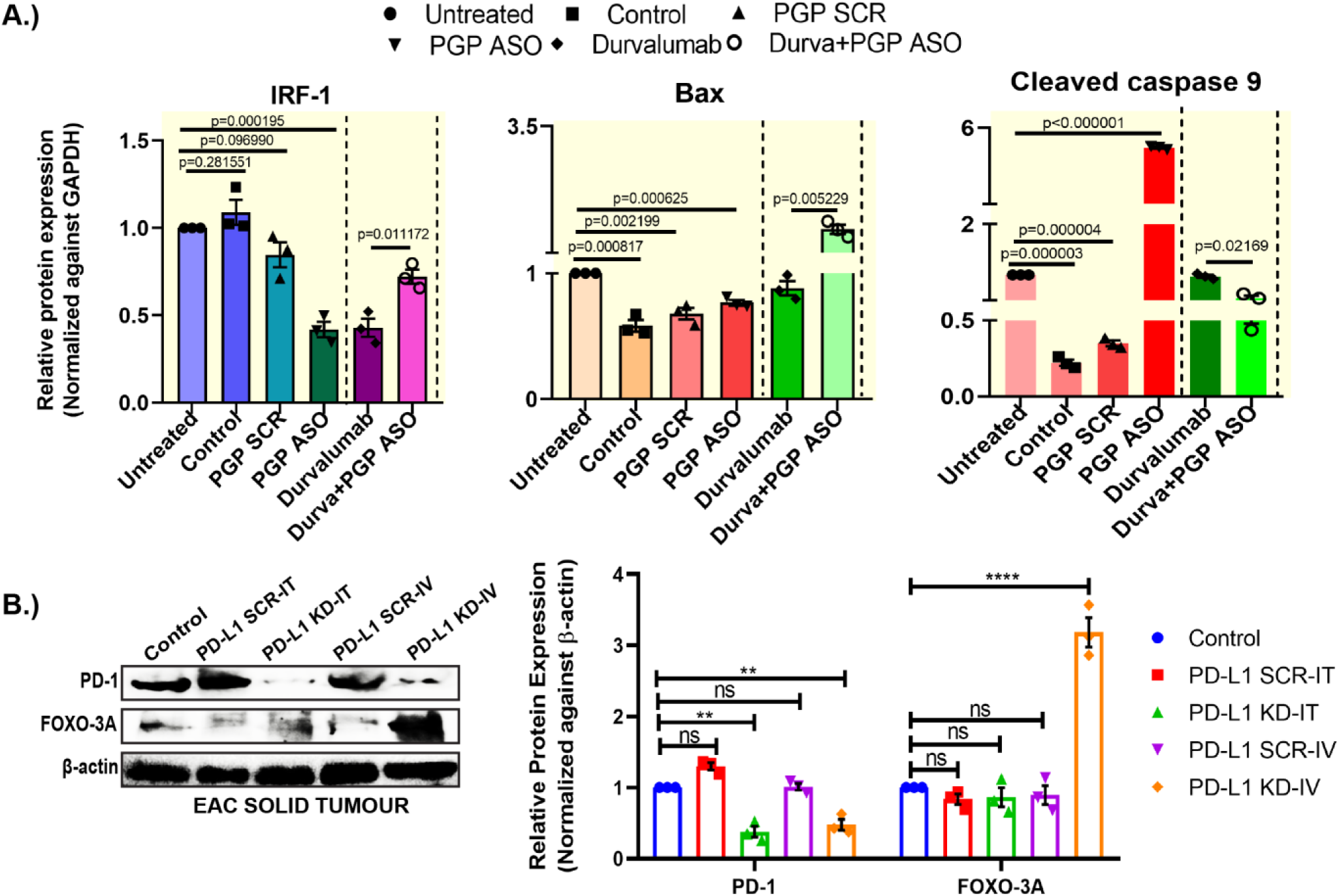
PGP ASO enhance the T cell function in *in vitro* and *in vivo*. **(A)** Graphical representation of densitometric analysis of IRF-1, Bax, cleaved caspase 9 proteins represented in figure 5B1. **(B)** Immunoblot analysis of PD-1 and FOXO-3A in PD-L1 KD mice (Left panel) and densitometric analysis of above said proteins (Right panel). Data shown as mean ± SEM, Ordinary Two-way ANOVA was performed followed by sidak multiple comparison test. \**p* ≤ 0.5 considered as statistically significant, ** and *** considered as higher significant than *p* ≤ 0.5.

**Figure S17:**
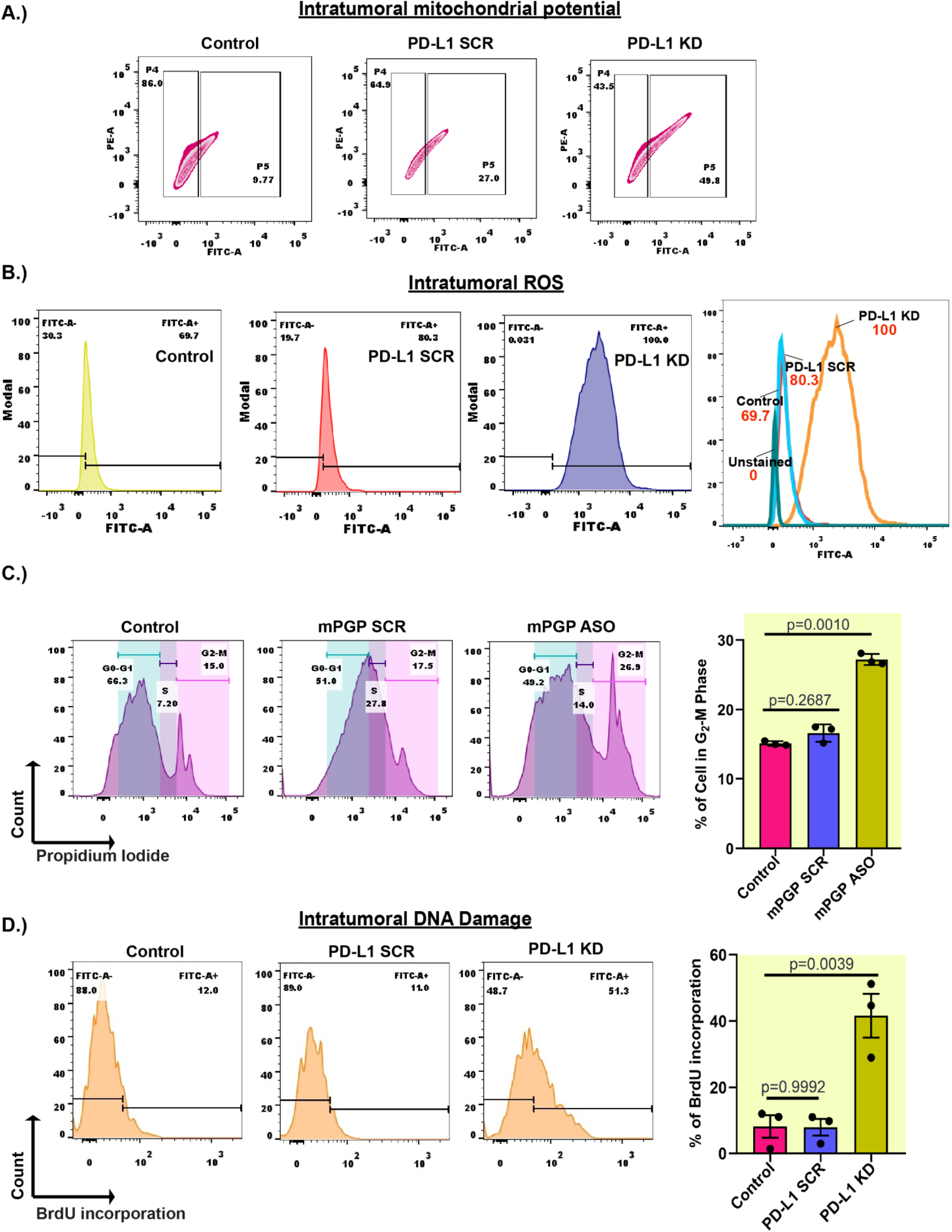
Tumour burden decreases in PD-L1 KD mice. **(A)** Depiction of intra-tumoral mitochondrial potential determination in PD-L1 KD mice. **(B)** Depiction of intra-tumoral ROS generation in PD-L1 KD mice. **(C)** Flow cytometric analysis of cell cycle arrest in EAC primary culture after mPGP ASO treatment. **(D**) Depiction of BrdU incorporation into tumour mass of PD-L1 KD mice. Data shown as means ± SEM, Ordinary one-way ANOVA was performed followed by sidak multiple comparison test. *p* ≤ 0.5 considered as statistically significant.

**Figure S18:**
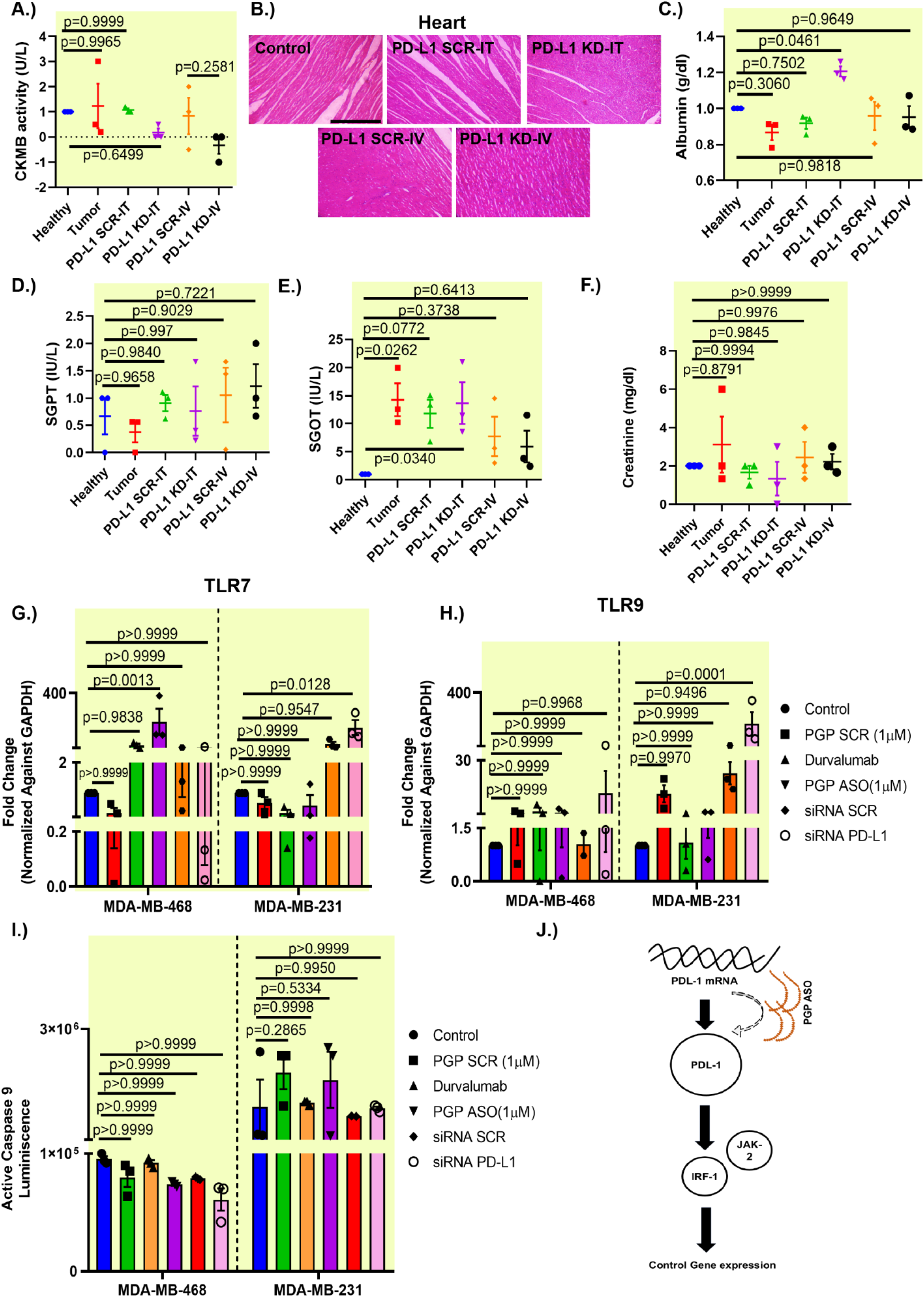
Off target effect evaluation of PGP ASO/mPGP ASO. **(A)** Assessment of serum CK-MB level in comparison with healthy (non-tumour bearing) mice. (B) H&E staining of cardiac tissue of PD-L1 SCR/KD mice, Scale bar= 400μm. Assessment of serum levels of (**C**) Albumin (**D**) SGPT (**E**) SGOT (**F**) creatinine of PD-L1 SCR/KD mice. mRNA expression patterns of (**G**)TLR7 (**H**) TLR9 in MDA-MB-231 and MDA-MB-468 cells. Data shown as mean ± SEM, Ordinary one-way ANOVA was performed followed by Dunnett’s multiple comparison test. *p* ≤ 0.5 considered as statistically significant. (**I**) Active Caspase glo 9 assessments in MDA-MB-231 and MDA-MB-468 cells. Data shown as means ± SEM, Ordinary one-way ANOVA was performed followed by sidak multiple comparison test. *p* ≤ 0.5 considered as statistically significant. (**J**) Diagram of PGP ASO mediated regulation of PD-L1 and downstream pathway.

**Figure S19:**
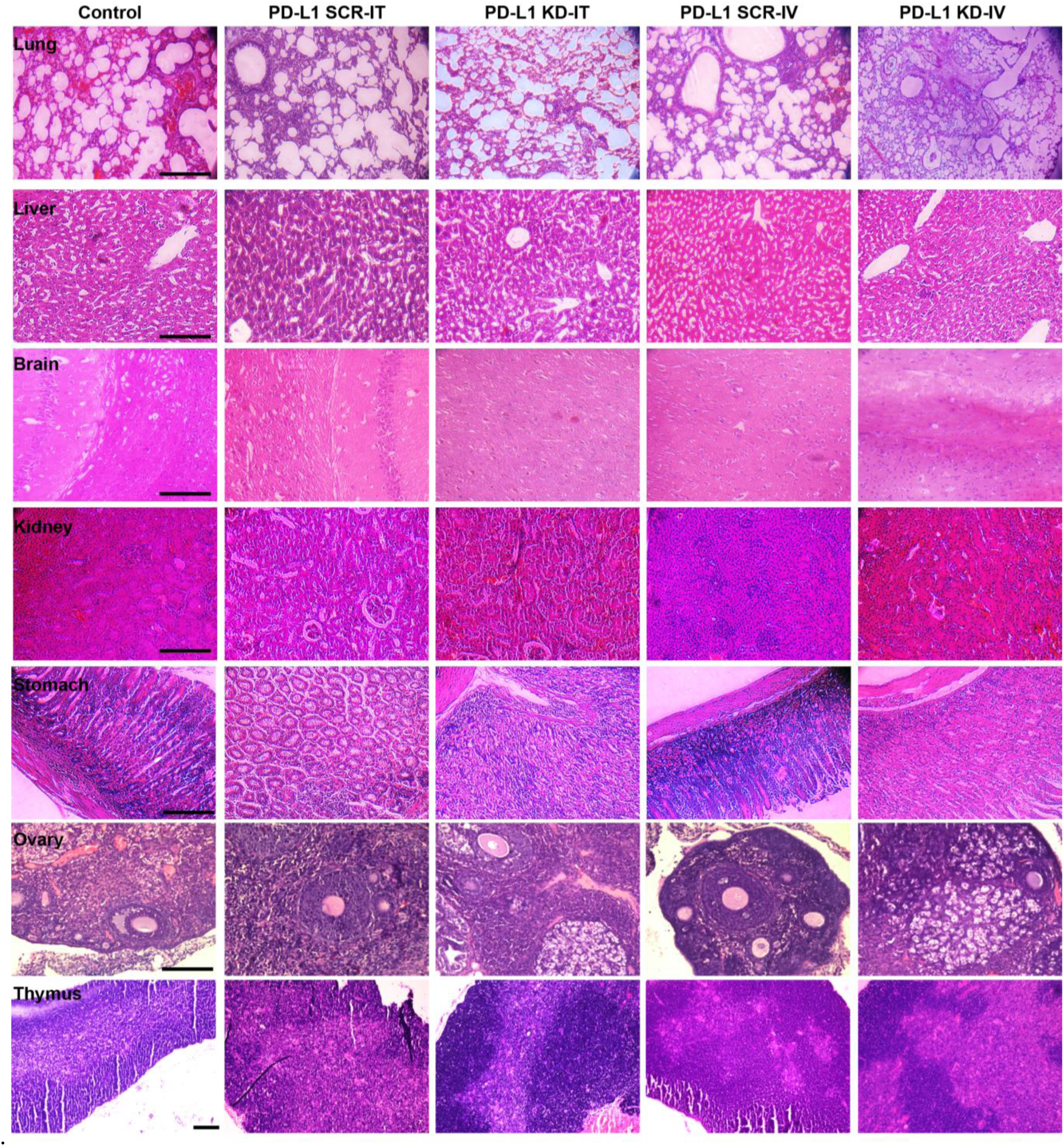
Off target effect evaluation in mPGP ASO treatment. H&E staining of Lung, liver, brain, kidney, stomach, ovary and thymus in PD-L1 SCR/KD mice.

**Table S1:**
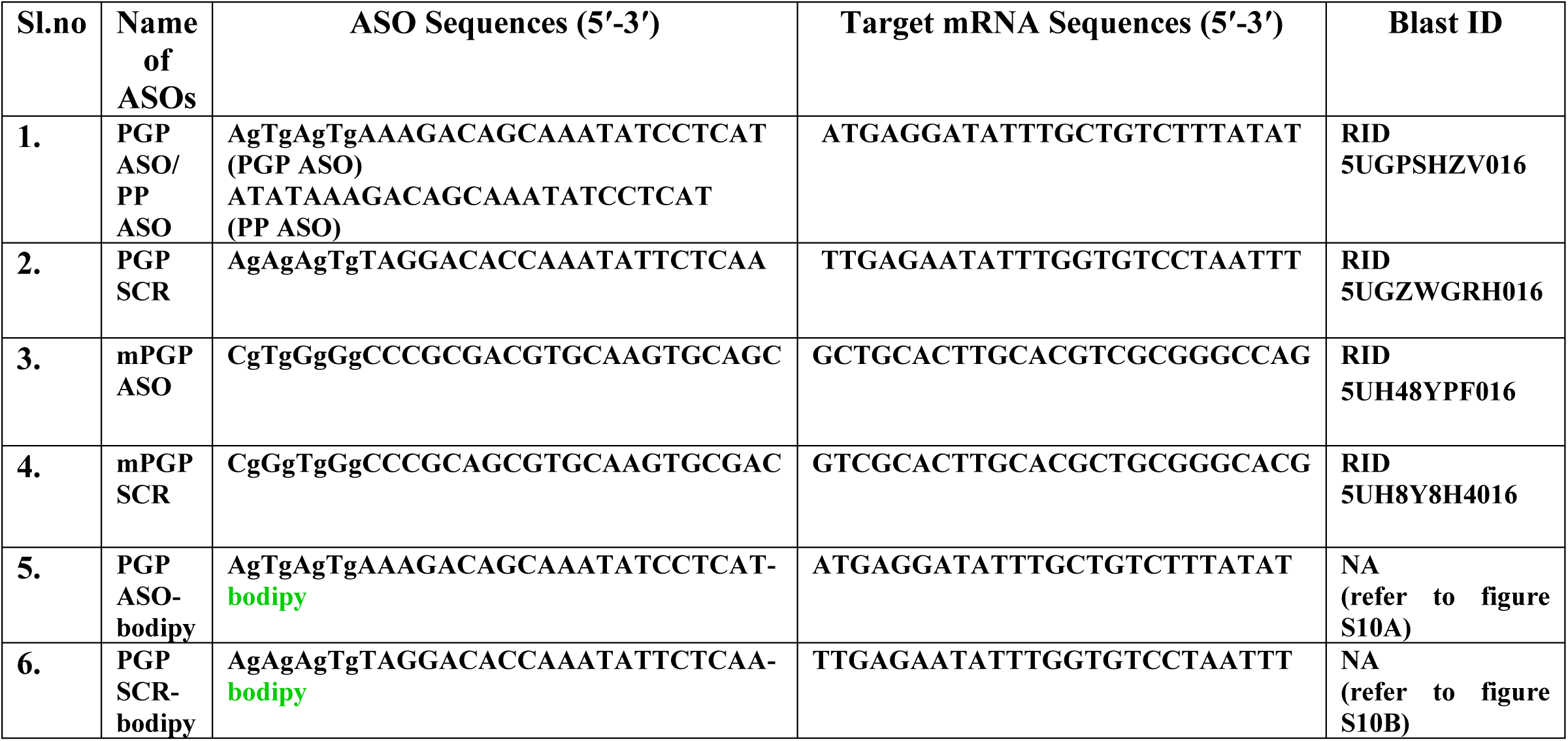
Nucleotide Sequences and corresponding Blast ID.

**Table S2:**
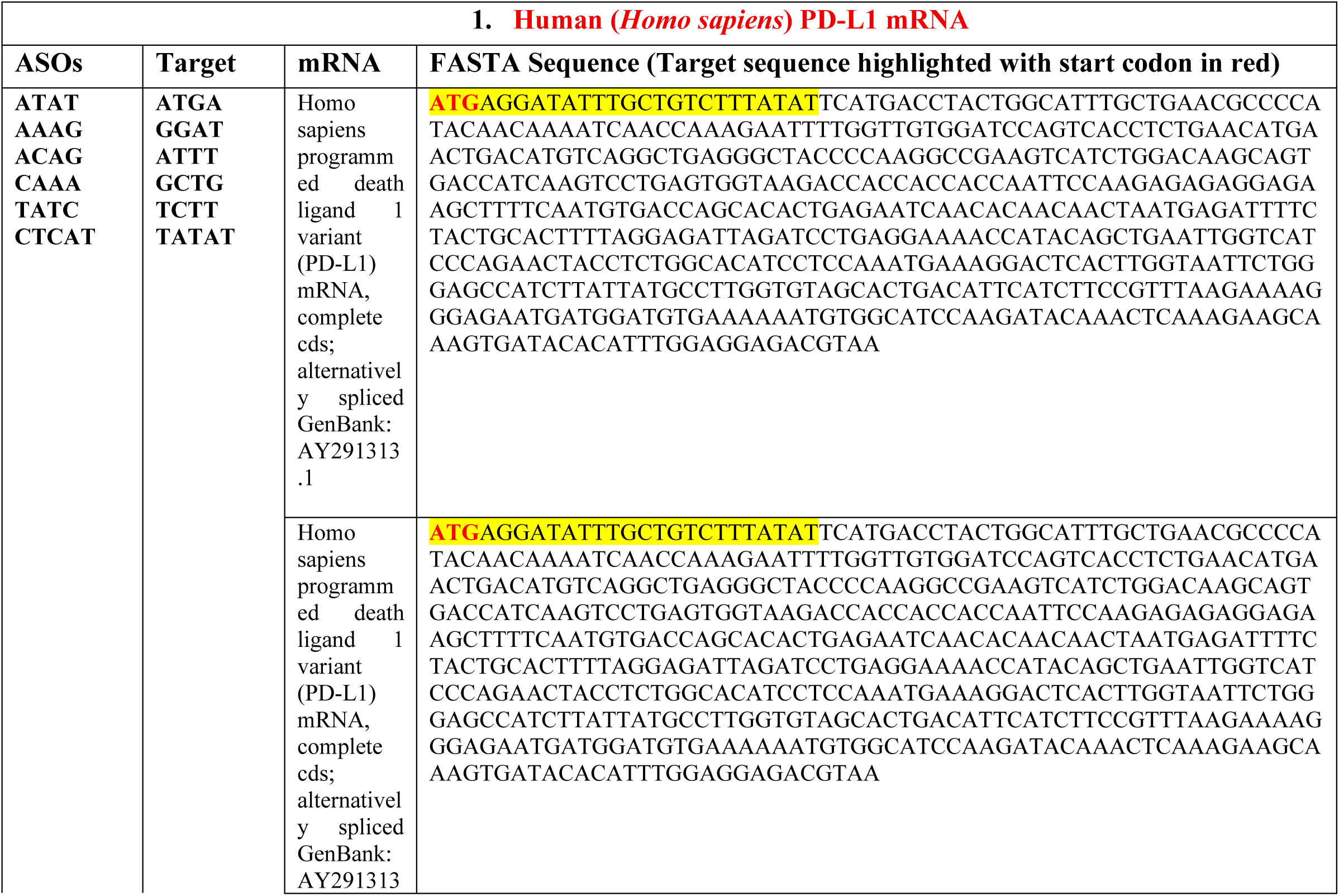
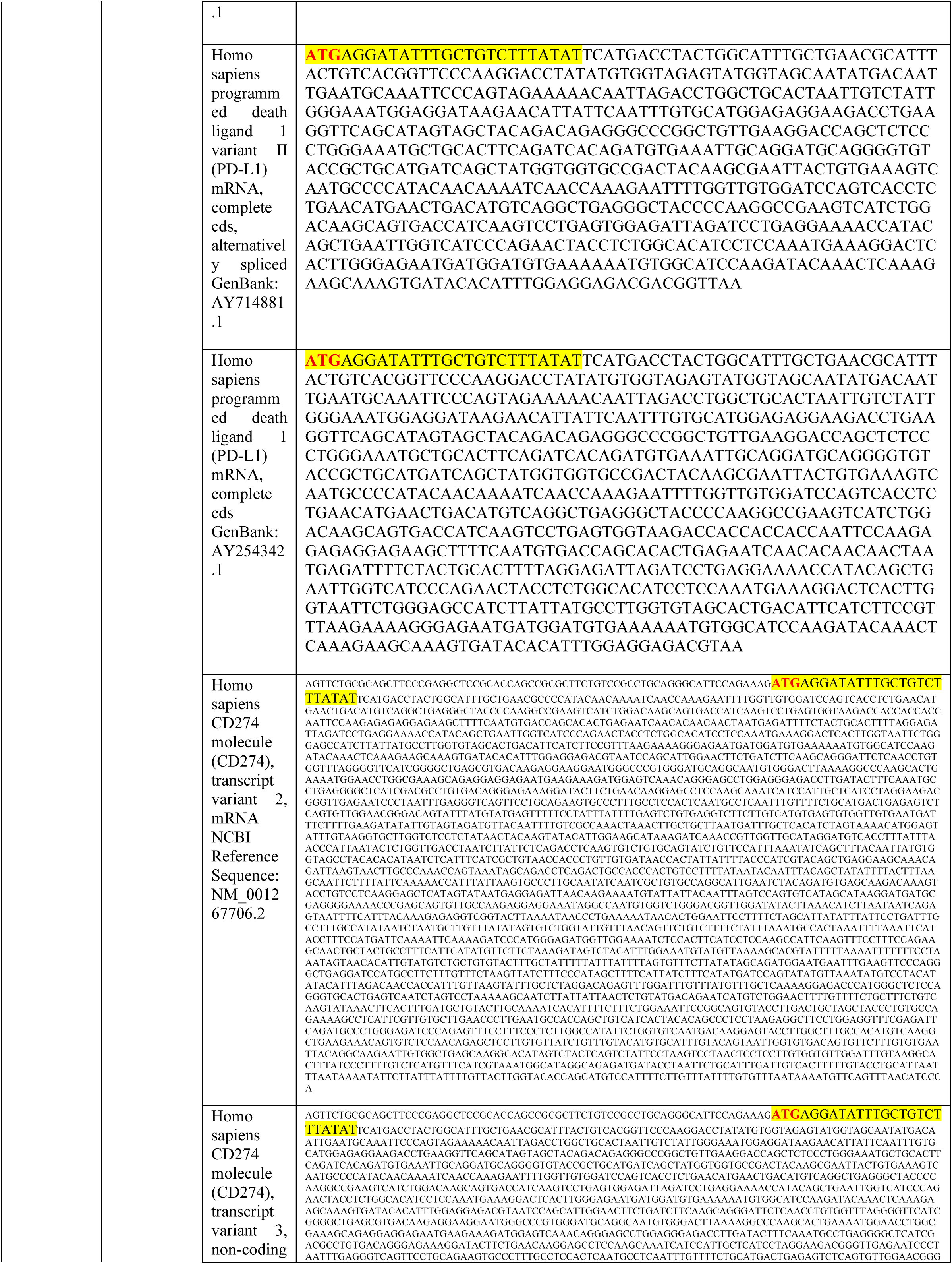
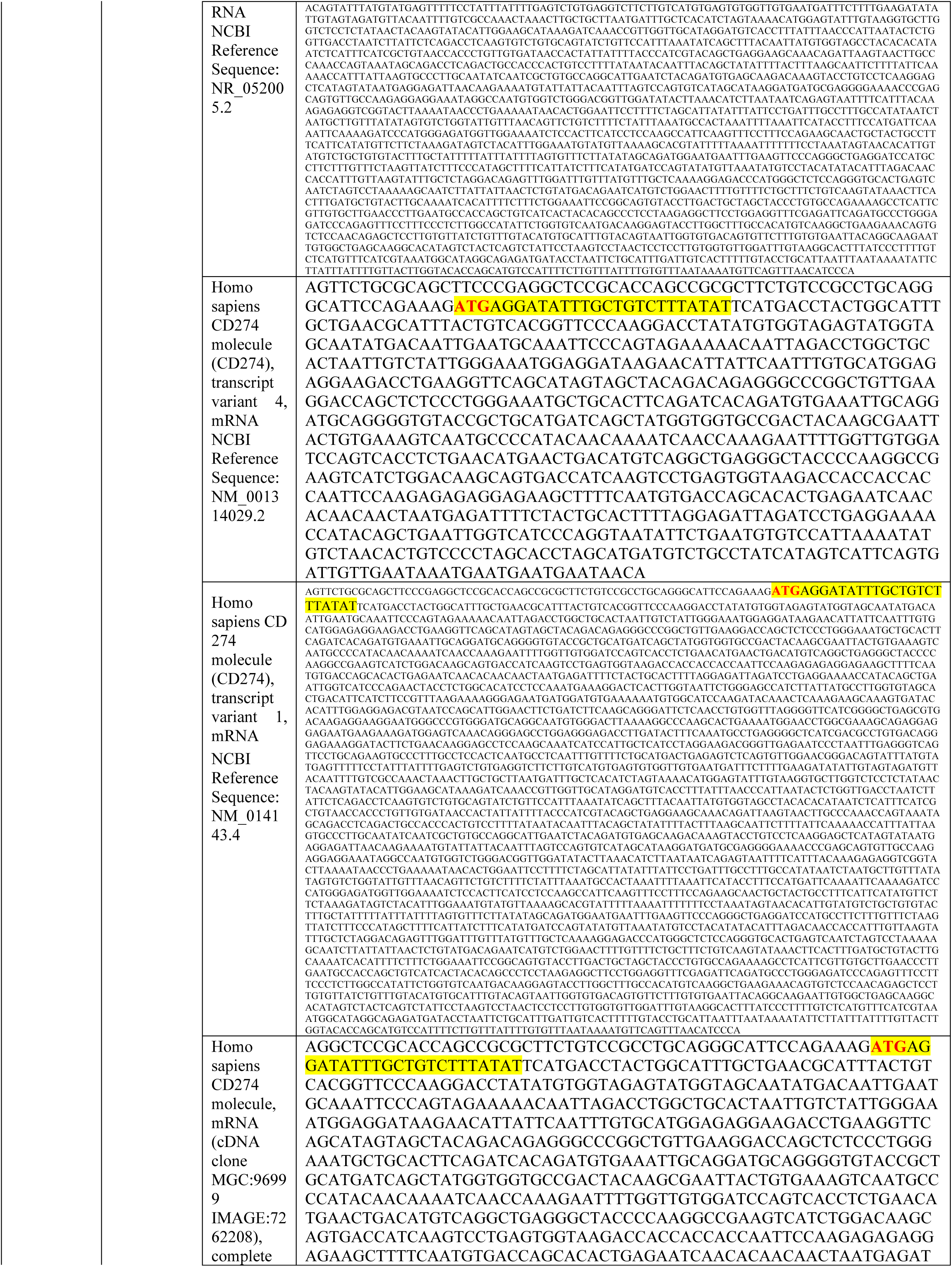
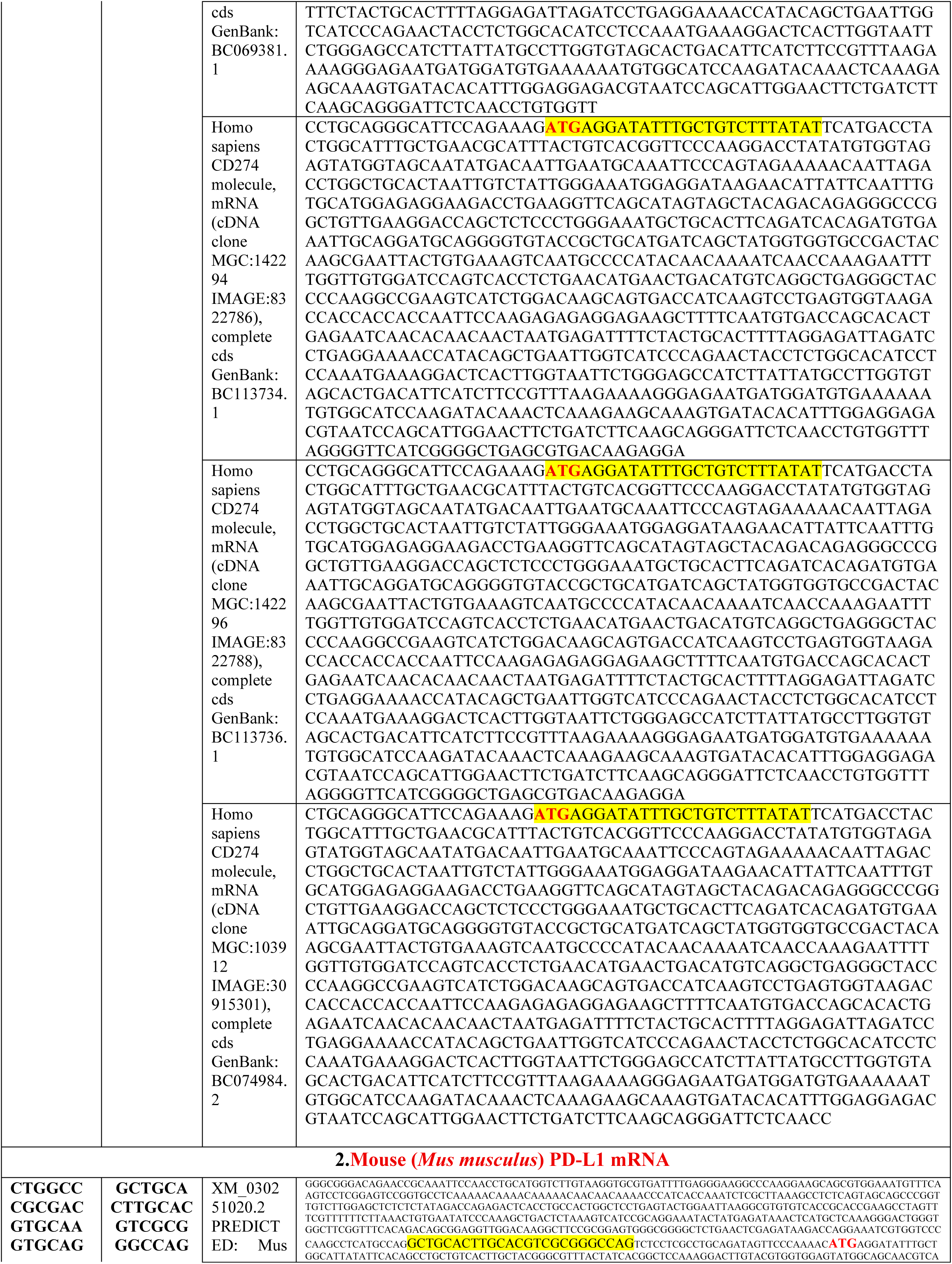
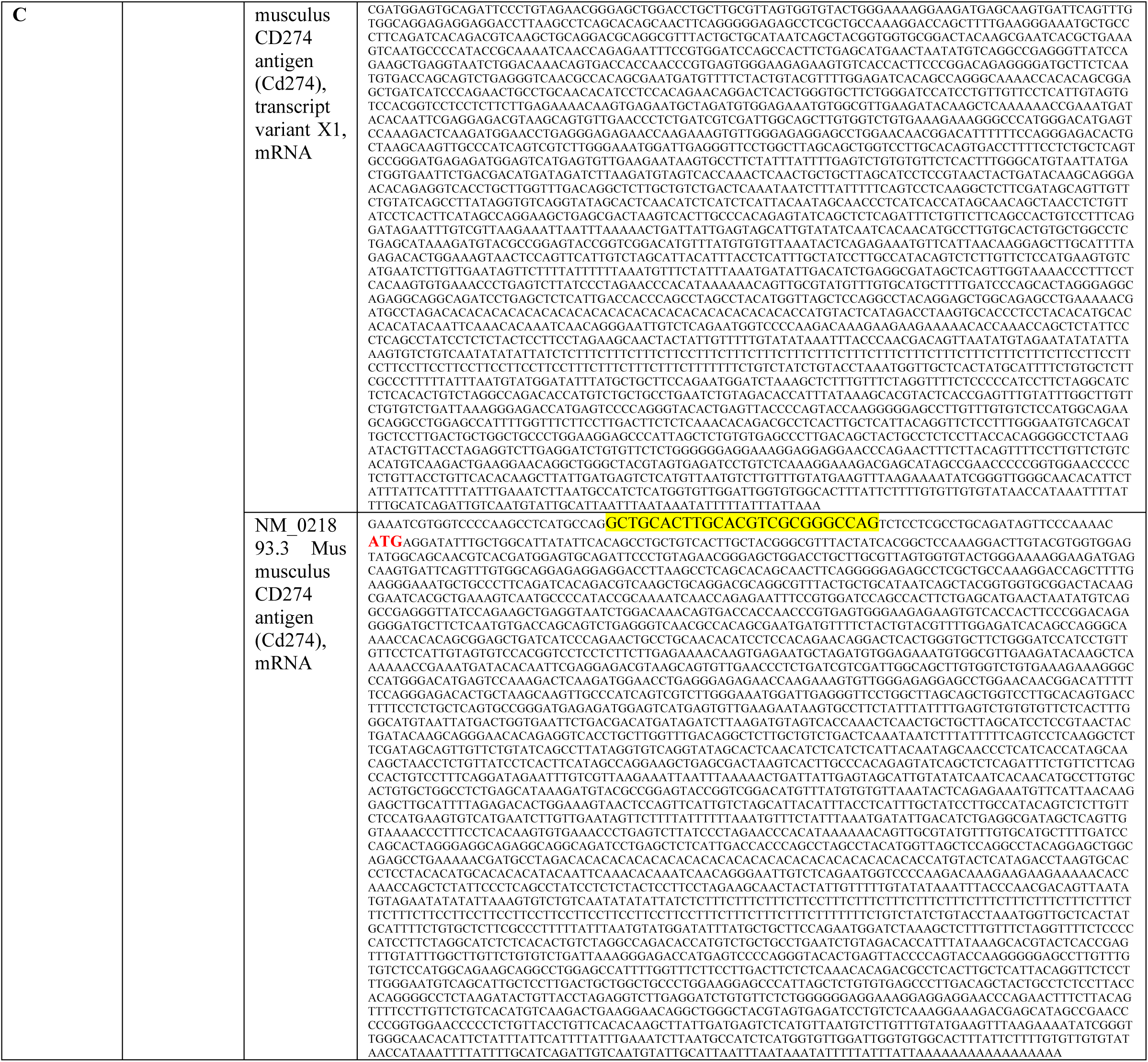
mRNA (including spliced variants) sequences of the targeted genes in FASTA format (target sequences of ASOs are highlighted and the start codons are marked in red).

**Table S3:**
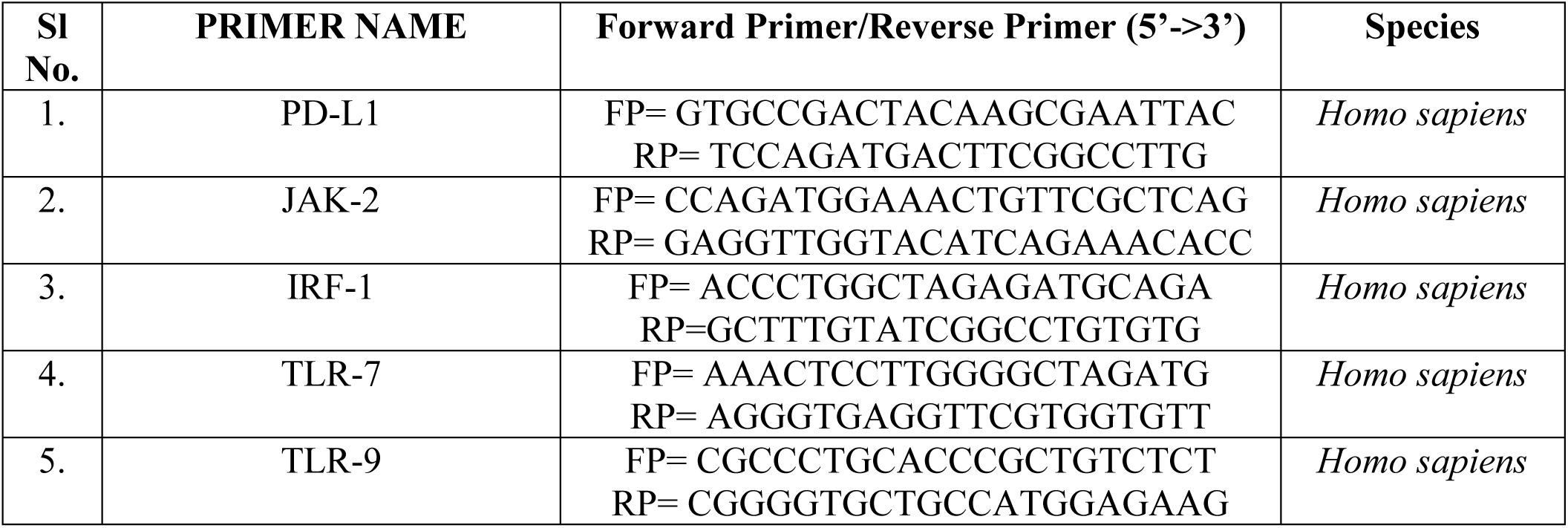
Primers used in PCR and RT-qPCR studies.

**Table S4:**
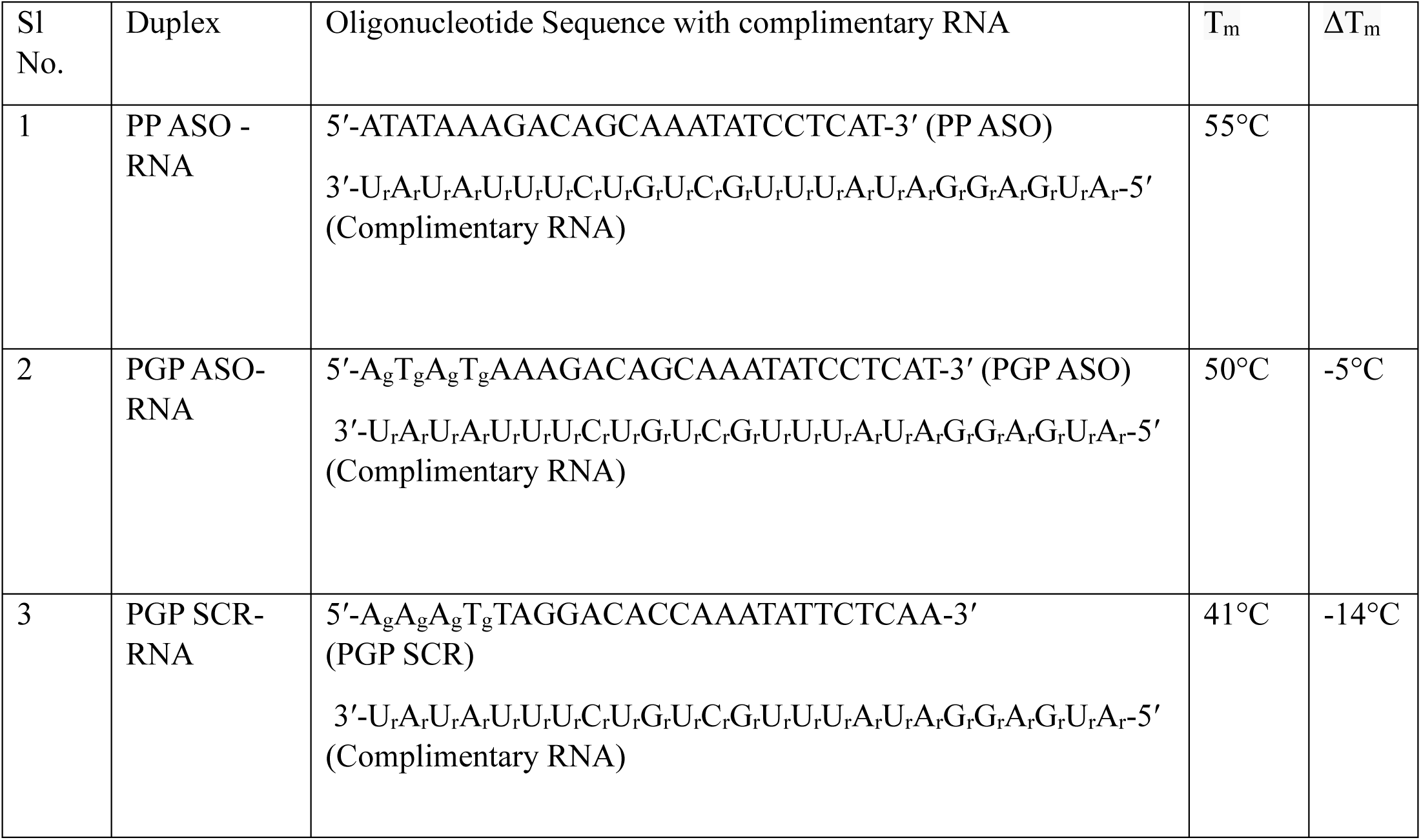
T_m_ data of duplexes with PMO and GMO-PMO chimera containing complementary RNA. (Conditions: 40 mM phosphate buffer (pH 7). The concentration of each strand was 1 μM. The T_m_ values reported are the averages of two independent experiments, and results differed by less than ±1.0°C. The ΔT_m_ values are in comparison to regular PMO-RNA).

## References

[1] Pardoll, Drew M. “The blockade of immune checkpoints in cancer immunotherapy.” Nat Rev Cancer 12, no. 4 (2012): 252–264.

[2] Doroshow, Deborah Blythe, Sheena Bhalla, Mary Beth Beasley, Lynette M. Sholl, Keith M. Kerr, Sacha Gnjatic, Ignacio I. Wistuba, David L. Rimm, Ming Sound Tsao, and Fred R. Hirsch. “PD-L1 as a biomarker of response to immune-checkpoint inhibitors.” Nat. Rev. Clin. Oncol. 18, no. 6 (2021): 345–362.

[3] Bardhan, Kankana, Theodora Anagnostou, and Vassiliki A. Boussiotis. “The PD1: PD-L1/2 pathway from discovery to clinical implementation.” Front. immunol. 7 (2016): 550.

[4] Zou, Weiping, Jedd D. Wolchok, and Lieping Chen. “PD-L1 (B7-H1) and PD-1 pathway blockade for cancer therapy: Mechanisms, response biomarkers, and combinations.” Sci. Transl. Med. 8, no. 328 (2016): 328rv4–328rv4.

[5] Sul, Joohee, Gideon M. Blumenthal, Xiaoping Jiang, Kun He, Patricia Keegan, and Richard Pazdur. “FDA approval summary: pembrolizumab for the treatment of patients with metastatic non-small cell lung cancer whose tumors express programmed death-ligand 1.” The oncologist 21, no. 5 (2016): 643–650.

[6] Lyons, Tomas G., Maura N. Dickler, and Elizabeth E. Comen. “Checkpoint inhibitors in the treatment of breast cancer.” Curr. Oncol. Rep. 20, no. 7 (2018): 51.

[7] Robert, Caroline, Jacob Schachter, Georgina V. Long, Ana Arance, Jean Jacques Grob, Laurent Mortier, Adil Daud et al. “Pembrolizumab versus ipilimumab in advanced melanoma.” NEJM 372, no. 26 (2015): 2521–2532.

[8] Gong, Jun, Alexander Chehrazi-Raffle, Srikanth Reddi, and Ravi Salgia. “Development of PD-1 and PD-L1 inhibitors as a form of cancer immunotherapy: a comprehensive review of registration trials and future considerations.” JITC 6, no. 1 (2018): 8.

[9] Lucibello, Giulia, Baharia Mograbi, Gerard Milano, Paul Hofman, and Patrick Brest. “PD-L1 regulation revisited: impact on immunotherapeutic strategies.” Trends Mol Med 27, no. 9 (2021): 868–881.

[10] Gao, Fan, Wei You, Lei Zhang, Ai-Zong Shen, Guang Chen, Ze Zhang, Xuan Nie et al. “Copper chelate targeting externalized phosphatidylserine inhibits PD-L1 expression and enhances Cancer immunotherapy.” JACS 147, no. 7 (2025): 5796–5807.

[11] Saha, Tanmoy, Michaela Fojtů, Astha Vinay Nagar, Liya Thurakkal, Balaaji Baanupriya Srinivasan, Meghma Mukherjee, Astralina Sibiyon et al. “Antibody nanoparticle conjugate–based targeted immunotherapy for non–small cell lung cancer.” Sci. Adv 10, no. 24 (2024): eadi2046.

[12] Li, Chia-Wei, Seung-Oe Lim, Weiya Xia, Heng-Huan Lee, Li-Chuan Chan, Chu-Wei Kuo, Kay-Hooi Khoo et al. “Glycosylation and stabilization of programmed death ligand-1 suppresses T-cell activity.” Nat. Commun. 7, no. 1 (2016): 12632.

[13] Escors, David, María Gato-Cañas, Miren Zuazo, Hugo Arasanz, María Jesus García-Granda, Ruth Vera, and Grazyna Kochan. “The intracellular signalosome of PD-L1 in cancer cells.” STTT 3, no. 1 (2018): 26.

[14] Gou, Qian, Chen Dong, Huihui Xu, Bibimaryam Khan, Jianhua Jin, Qian Liu, Juanjuan Shi, and Yongzhong Hou. “PD-L1 degradation pathway and immunotherapy for cancer.” Cell Death Dis 11, no. 11 (2020): 955.

[15] Dume, Bogdan, Emilia Licarete, and Manuela Banciu. “Advancing cancer treatments: The role of oligonucleotide-based therapies in driving progress.” Mol. Ther. Nucleic Acids 35, no. 3 (2024).

[16] Duan, Dongsheng, Nathalie Goemans, Shin’ichi Takeda, Eugenio Mercuri, and Annemieke Aartsma-Rus. “Duchenne muscular dystrophy.” Nat. Rev. Dis. Primers 7, no. 1 (2021): 13.

[17] Tarbashevich, Katsiaryna, Atanu Ghosh, Arnab Das, Debajyoti Kuilya, Swrajit Nath Sharma, Surajit Sinha, and Erez Raz. “Optochemical control over mRNA translation by photocaged phosphorodiamidate morpholino oligonucleotides in vivo.” Nat. Commun. 16, no. 1 (2025): 3614.

[18] Kundu, Jayanta, Atanu Ghosh, Ujjwal Ghosh, Arnab Das, Dhriti Nagar, Sankha Pattanayak, Aurnab Ghose, and Surajit Sinha. “Synthesis of phosphorodiamidate morpholino oligonucleotides using trityl and fmoc chemistry in an automated oligo synthesizer.” J. Org. Chem 87, no. 15 (2022): 9466–9478.

[19] Foulkes, William D., Ian E. Smith, and Jorge S. Reis-Filho. “Triple-negative breast cancer.” NEJM 363, no. 20 (2010): 1938–1948.

[20] Mandapati, Aditya, and Kiven Erique Lukong. “Triple negative breast cancer: approved treatment options and their mechanisms of action.” J Cancer Res Clin Oncol 149, no. 7 (2023): 3701–3719.

[21] Dent, Rebecca, Wedad M. Hanna, Maureen Trudeau, Ellen Rawlinson, Ping Sun, and Steven A. Narod. “Pattern of metastatic spread in triple-negative breast cancer.” Breast Cancer Res Treat 115, no. 2 (2009): 423–428.

[22] Das, Ujjal, Jayanta Kundu, Pallab Shaw, Chandra Bose, Atanu Ghosh, Shalini Gupta, Sudipta Sarkar, Jhuma Bhadra, and Surajit Sinha. “Self-transfecting GMO-PMO chimera targeting Nanog enable gene silencing in vitro and suppresses tumor growth in 4T1 allografts in mouse.” Mol. Ther. Nucleic Acids 32 (2023): 203–228.

[23] Mishra, Sneha, Ankit Kumar Tamta, Mohsen Sarikhani, Perumal Arumugam Desingu, Shruti M. Kizkekra, Anwit Shriniwas Pandit, Shweta Kumar, Danish Khan, Sathees C. Raghavan, and Nagalingam R. Sundaresan. “Subcutaneous Ehrlich Ascites Carcinoma mice model for studying cancer-induced cardiomyopathy.” Sci. Rep. 8, no. 1 (2018): 5599.

[24] Lin, Xianbin, Liangan Lin, Jingyang Wu, Wentan Jiang, Jiayun Wu, Jianshen Yang, and Chun Chen. “A targeted siRNA-loaded PDL1-exosome and functional evaluation against lung cancer.” “Thorac. Cancer” 13, no. 11 (2022): 1691–1702.

[25] Pacheco-Torres, Jesus, Marie-France Penet, Balaji Krishnamachary, Yelena Mironchik, Zhihang Chen, and Zaver M. Bhujwalla. “PD-L1 siRNA theranostics with a dextran nanoparticle highlights the importance of nanoparticle delivery for effective tumor PD-L1 downregulation.” Front. Oncol 10 (2021): 614365.

[26] Luo, Fatao, Gang Yang, Xia Bai, Deyu Yuan, Ling Li, Diyue Wang, Xiaoxiang Lu et al. “Anti-tumor effect of PD-L1-targeting antagonistic aptamer-ASO delivery system with dual inhibitory function in immunotherapy.” Cell Chem. Biol 30, no. 11 (2023): 1390–1401.

[27] Ghosh, Atanu, Arpan Banerjee, Shalini Gupta, and Surajit Sinha. “A unified phosphoramidite platform for the synthesis of morpholino oligonucleotides and diverse chimeric backbones.” JACS 146, no. 48 (2024): 32989–33001.

[28] Jorgovanovic, Dragica, Mengjia Song, Liping Wang, and Yi Zhang. “Roles of IFN-γ in tumor progression and regression: a review.” Biomark Res. 8, no. 1 (2020): 49.

[29] Mauthe, Mario, Idil Orhon, Cecilia Rocchi, Xingdong Zhou, Morten Luhr, Kerst-Jan Hijlkema, Robert P. Coppes, Nikolai Engedal, Muriel Mari, and Fulvio Reggiori. “Chloroquine inhibits autophagic flux by decreasing autophagosome-lysosome fusion.” Autophagy 14, no. 8 (2018): 1435–1455.

[30] Ye, Hongxing, Mantao Chen, Fei Cao, Hongguang Huang, Renya Zhan, and Xiujue Zheng. “Chloroquine, an autophagy inhibitor, potentiates the radiosensitivity of glioma initiating cells by inhibiting autophagy and activating apoptosis.” BMC neurology 16, no. 1 (2016): 178.

[31] Feng, Chong, Lening Zhang, Xin Chang, Dongliang Qin, and Tao Zhang. “Regulation of post-translational modification of PD-L1 and advances in tumor immunotherapy.” Front. immunol. 14 (2023): 1230135.

[32] Alvarez-Argote, Juliana, and Constantin A. Dasanu. “Durvalumab in cancer medicine: a comprehensive review.” Expert Opin. Biol. Ther. 19, no. 9 (2019): 927–935.

[33] Gao, Yang, Naoe Taira Nihira, Xia Bu, Chen Chu, Jinfang Zhang, Aleksandra Kolodziejczyk, Yizeng Fan et al. “Acetylation-dependent regulation of PD-L1 nuclear translocation dictates the efficacy of anti-PD-1 immunotherapy.” Nat. Cell Biol 22, no. 9 (2020): 1064–1075.

[34] Sun, Hao, Yingmei Li, Peng Zhang, Haizhou Xing, Song Zhao, Yongping Song, Dingming Wan, and Jifeng Yu. “Targeting toll-like receptor 7/8 for immunotherapy: recent advances and prospectives.” Biomark Res. 10, no. 1 (2022): 89.

[35] Wang, Zhisong, Yan Gao, Lei He, Shuhao Sun, Tingting Xia, Lu Hu, Licheng Yao et al. “Structure-based design of highly potent toll-like receptor 7/8 dual agonists for cancer immunotherapy.” J. Med. Chem. 64, no. 11 (2021): 7507–7532.

[36] Goyenvalle, Aurélie, Cecilia Jimenez-Mallebrera, Willeke van Roon, Sabine Sewing, Arthur M. Krieg, Virginia Arechavala-Gomeza, and Patrik Andersson. “Considerations in the preclinical assessment of the safety of antisense oligonucleotides.” Nucleic Acid Ther. 33, no. 1 (2023): 1–16.

[37] Lopez, Diana M., Vijaya Charyulu, and Becky Adkins. “Influence of breast cancer on thymic function in mice.” J Mammary Gland Biol Neoplasia 7, no. 2 (2002): 191–199.

[38] Wu, Bo, Bo Zhang, Bowen Li, Haoqi Wu, and Meixi Jiang. “Cold and hot tumors: from molecular mechanisms to targeted therapy.” Signal Transduct. Target. Ther. 9, no. 1 (2024): 274.

[39] Garcia-Diaz, Angel, Daniel Sanghoon Shin, Blanca Homet Moreno, Justin Saco, Helena Escuin-Ordinas, Gabriel Abril Rodriguez, Jesse M. Zaretsky et al. “Interferon receptor signaling pathways regulating PD-L1 and PD-L2 expression.” Cell Rep. 19, no. 6 (2017): 1189–1201.

[40] Mimura, Kousaku, Jun Liang Teh, Hirokazu Okayama, Kensuke Shiraishi, Ley-Fang Kua, Vivien Koh, Duane T. Smoot et al. “PD-L1 expression is mainly regulated by interferon gamma associated with JAK-STAT pathway in gastric cancer.” Cancer Sci. 109, no. 1 (2018): 43–53.

[41] Lemma, Eyoel Yemanaberhan, Anudari Letian, Nasser K. Altorki, and Timothy E. McGraw. “Regulation of PD-L1 trafficking from synthesis to degradation.” Cancer Immunol. Res. 11, no. 7 (2023): 866–874.

[42] Bertrand, Florie, Anne Montfort, Elie Marcheteau, Caroline Imbert, Julia Gilhodes, Thomas Filleron, Philippe Rochaix et al. “TNFα blockade overcomes resistance to anti-PD-1 in experimental melanoma.” Nat. Commun. 8, no. 1 (2017): 2256.

[43] Rahat, Michal A., and Bernhard Hemmerlein. “Macrophage-tumor cell interactions regulate the function of nitric oxide.” Front. Physiol. 4 (2013): 144.

[44] Churlaud, Guillaume, Fabien Pitoiset, Fadi Jebbawi, Roberta Lorenzon, Bertrand Bellier, Michelle Rosenzwajg, and David Klatzmann. “Human and mouse CD8+ CD25+ FOXP3+ regulatory T cells at steady state and during interleukin-2 therapy.” Front. immunol. 6 (2015): 171.

[45] Hariyanto, Agustinus Darmadi, Tiara Bunga Mayang Permata, and Soehartati Argadikoesoema Gondhowiardjo. “Role of CD4+ CD25+ FOXP3+ TReg cells on tumor immunity.” Immunol Med. 45, no. 2 (2022): 94–107.

[46] Moulton, Jon D., and Shan Jiang. “Gene knockdowns in adult animals: PPMOs and vivo-morpholinos.” Molecules 14, no. 3 (2009): 1304–1323.

[47] Pan, Simin, Michael Cesarek, Carla Godoy, Cynthia M. Co, Catherine Schindler, Kelbi Padilla, Andrew Haskell et al. “Morpholino-driven blockade of Dkk-1 in osteosarcoma inhibits bone damage and tumour expansion by multiple mechanisms.” Br J Cancer. 127, no. 1 (2022): 43–55.

[48] Ferguson, David P., Lawrence J. Dangott, and J. Timothy Lightfoot. “Lessons learned from vivo-morpholinos: How to avoid vivo-morpholino toxicity.” Biotechniques 56, no. 5 (2014): 251–256.

